# Sequential Activation of Guide RNAs for Algorithmic Multiplexing of Cas9 Activities

**DOI:** 10.1101/2020.06.20.162982

**Authors:** Ryan Clarke, Alexander R. Terry, Hannah Pennington, Matthew S. MacDougall, Maureen Regan, Bradley J. Merrill

## Abstract

Genetic manipulation of mammalian cells is instrumental to modern biomedical research but is currently limited by poor capabilities of sequentially controlling multiple manipulations in cells. Currently, either highly multiplexed manipulations can be delivered to populations of cells all at one time, or gene regulatory sequences can be engineered to conditionally activate a few manipulations within individual cells. Here, we provide proof-of-principle for a new system enabling multiple genetic manipulations to be executed as a preprogrammed cascade of events. The system leverages the programmability of the *S. pyogenes* Cas9 RNA-guided nuclease and is based on flexible arrangements of individual modules of activity. The basic module consists of an inactive single guide RNA (sgRNA) - like component that is converted to an active state through the effects of another sgRNA. Modules can be arranged to bring about an algorithmic program of genetic manipulations without the need for engineering cell type specific promoters or gene regulatory sequences. With the expanding diversity of available tools that utilize spCas9 to edit, repress or activate genes, this sgRNA-based system provides multiple levels for interfacing with host cell biology. In addition, ability of the system to progress through multiple modules from episomal plasmid DNA makes it suitable for applications sensitive to the presence of heterologous genomic DNA sequences and broadly applicable to biomedical research and mammalian cell engineering.

## INTRODUCTION

RNA-guided nucleases have empowered manipulation of mammalian genes for research and therapies at diverse scales, from individual cells to whole organs. Although other nucleases, including synthetic proteins designed to target specific DNA sequences (i.e. zinc finger nucleases, TALENs), remain effective genome editing reagents, RNA-guided nucleases benefit from the simplicity by which they can be programmed to target a DNA sequence, which is accomplished by Watson-Crick base pairing of a portion of a guide RNA to its DNA target (Cong et al., 2013; Jinek et al., 2012; Jinek et al., 2013; Mali et al., 2013). The ease of changing the targeting sequence and the capability of targeting virtually any DNA sequence has stimulated an explosion in the usage of RNA-guided nucleases as the predominant gene-editing tool.

The spCas9 nuclease used for most genome editing in mammals was derived from the clustered regularly interspaced palindromic repeats (CRISPR)-Cas9 system in *S. pyogenes* (Gasiunas et al., 2012; Jinek et al., 2012). It targets DNA with the so-called “spacer” sequence, which is the first 20nt of a single guide RNA (sgRNA), and it requires an NGG motif to be adjacent to the spacer sequence for nuclease activity (Gasiunas et al., 2012; Jinek et al., 2012). These properties of spCas9 have made it deployable in highly multiplexed modes; libraries of sgRNA can direct spCas9 to mutate hundreds of thousands of sites in large populations of cells (Shalem et al., 2014; Wang et al., 2014) or even thousands of sites in individual cells (Smith et al., 2020).

In addition to genome editing, spCas9 has also been used for synthetic biology purposes, such as in generating genetic circuits that change cellular properties (Schwarz et al., 2017) and recording information about a cell in the cell’s genomic DNA sequence (Chan et al., 2019; Kalhor et al., 2018; Kalhor et al., 2017; McKenna et al., 2016; Perli et al., 2016; Schwarz et al., 2017). Whereas the flexibility in targeting DNA sequences is very useful for applying spCas9 for synthetic biology, the ability of spCas9 protein to simultaneously work with every functional sgRNA in a cell poses a barrier to constructing circuits that need to sequentially or temporally trigger independent sgRNA-mediated effects. This is a problem for genetic recording circuits, because spCas9 mutates all the target sequences at the same time, saturating the recording medium within a single or a few cell divisions. Some methods for prolonging the duration of spCas9 activity have been developed to expand the length of recording, either by reducing its activity (Chan et al., 2019; McKenna et al., 2016) or through self-targeting sgRNA, which encode iterative editing of their own sequence (Kalhor et al., 2017; Perli et al., 2016). A third approach is exemplified by the DOMINO system (Farzadfard et al., 2019), which demonstrates the concept of iterative genome edits to the spacer-target sequence pairs using base editors. By progressing through a series of states in a preprogrammed, stepwise manner, DOMINO added a critical ability to combine multiplexed activities as logical operations within a larger architecture (Farzadfard et al., 2019).

The use of sgRNA that sequentially regulate one another to mediate progression through successive states enables design of powerful systems for storing information in genomic DNA. However, the current approaches come with a tradeoff of limiting the ability of interfacing with the host genome; essentially, systems like DOMINO and self-targeting sgRNA need to use the spacer sequence of the sgRNA to power progression through iterative steps in a circuit. Because the sgRNA in these systems cannot be easily targeted to a variety of genomic DNA sequences, their capacity for controlling genome-encoded cellular activities is limited.

Currently, controlling a sequential order of sgRNA targeting various genomic sites requires exogenous inputs and is limited to two events. Regulating the stability of sgRNA via chemically-controlled ligands allows for certain species of sgRNA within a pool to be specifically controlled (Kundert et al., 2019; Tang et al., 2017). The expression and stability of multiple Cas9 orthologues can also be employed to regulate the order of gene targeting (Kim et al., 2019). Although these systems enable sequential Cas9 activities, they are limited to scaling based on the available small-molecule regulated aptamers, chemically-inducible degrons, and inducible promoters. Furthermore, consistent exogenous intervention to control steps within a program ignores the asynchronicity of cell populations and can reduce the frequency of properly responding cells as each step is initiated.

We sought to develop an open system that provides step-wise multiplexing and also liberates the spacer sequence for use in targeting spCas9 to the genome. By creating self-destructing sgRNA with the insertion of *cis*-cleaving ribozyme sequence, we show that sgRNA can be held in an inactive state until another distinct sgRNA converts them to become active by removal of the ribozyme sequence. These conditional sgRNA can be combined to form daisy-chain or ramified cascades to target spCas9 activities throughout the genome. Proof of concept experiments demonstrated functionality of four sequential steps in a daisy chain arrangement and cascades with two ramified branch points inducing two sequential genomic-encoded phenotypic changes in cells. We posit that application of the system will enable long-term genetic recording and programming of complex cellular biology in mammals.

## RESULTS

### The pro-Guide: an inactive sgRNA that can be converted to an active, mature-Guide state via genome editing

Inspired by synthetic biology in prokaryotes (Farzadfard et al., 2019; Tang and Liu, 2018), we reasoned that an ideal mammalian system would have capabilities of both seamlessly interfacing with the mammalian genome and progressing through a series of instructions independently of the cell’s state. To accomplish this, sgRNA were engineered to exist in an “off” inactive state until converted to the “on” active state via the genome editing function of a different sgRNA and spCas9 (Fig. 1A,B). We named these inactive sgRNA-like products “pro-Guides” (pGuide), because they are conceptually similar to prodrugs, which get converted to an active state by specific biochemistry they encounter in the body. Removal or disruption of the inactivating sequence converts a pGuide to a “mature-Guide” (mGuide), which has sgRNA-like activity (Fig. 1C). It was important that the pGuide-to-mGuide conversion was mediated by spCas9, because it provided a mechanism for arranging the multiple activities such that conversion of one pGuide could result in the conversion of another downstream pGuide (Fig. 1C). A simple daisy chain arrangement of pGuides, with each one activating the next and targeting a genomic site, would enable sequential editing of multiple genomic sites to occur via a preconfigured order of events (Fig. 1D).

**Figure 1:**
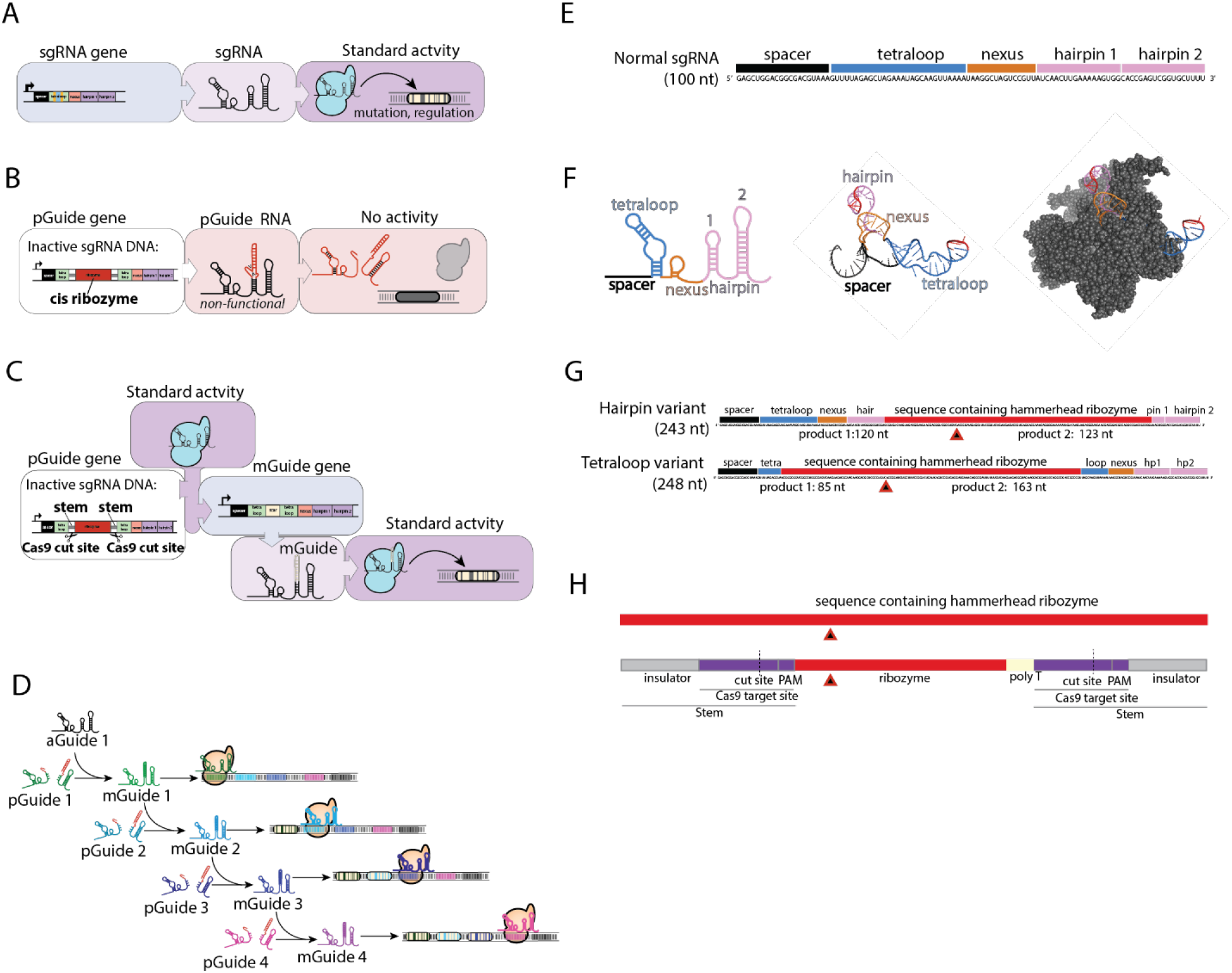
Using activatable sgRNA to program sequentiality into multiplexed Cas9 activities. A. A sgRNA gene encodes the target of spCas9 activity actuated by transcription of the sgRNA. Upon binding the target DNA sequence, a frequent result of spCas9-generated double stranded DNA breaks is a mutation at the target site. B. Regulation of spCas9 activity is achieved through generation of sgRNA-like molecules that are transcribed yet not suitable for functional spCas9 binding and subsequent activity. Through insertion of a *cis* cleaving ribozyme sequence (red) into the gene encoding the sgRNA, the RNA is cleaved upon ribozyme folding when the gene is transcribed. These “pro-guide RNA” or “pGuide” molecules are thus encoded to self-destruct until the gene is converted to an active state. Ribozyme encoding DNA sequences are shown inserted into hairpin 1 region of the sgRNA. C. Conversion of the inactive pGuide to form encoding the essential activity of an sgRNA (mature Guide or mGuide) is accomplished by spCas9-mediated genome editing. A gene edit removes the ribozyme encoding sequence from the pGuide generating an mGuide gene, which encodes an intact and active sgRNA-like mGuide RNA. The mGuide is functional for spCas9 binding and activity as a normal sgRNA; shown is an activity directing editing of a target sequence as illustrated for the sgRNA in (A). Note, spCas9 activity that inactivates the ribozyme can be introduced into a cell from an external source, or it could be derived following conversion of a pGuide to an mGuide gene already in the cell. The programmability of targeting spCas9 to various DNA sequences with the first 20nt of an sgRNA or mGuide enables conversion of multiple pGuide genes to be linked in preconfigured array, such as the one shown in (C). D. Schematic displaying sequentially programmable spCas9 activity via a linear cascade of pGuides. The cascade is designed so that the target sequence for an mGuide matches DNA sequences flanking the ribozyme in the next pGuide in a series of four pGuides. Prior to conversion of the first pGuide, the ribozyme in each pGuide inactivates them, preventing spCas9 activity at any of the potential target sites. The addition of an activating sgRNA (activating Guide or aGuide) targeting pGuide 1 initiates the linear cascade by converting the pGuide1 gene to an mGuide state. The mGuide1 RNA is designed to target pGuide 2 gene and convert it to an mGuide state. Continuation of the process for pGuide 3 and pGuide 4 illustrates activity at four different genomic target sites and their conversion to a mutated sequence through a stepwise process. E. Primary structure, nucleotide sequence, and arrangement of the *S. pyogenes* sgRNA motifs, which are described in panel (D). F. Left: secondary structure of the sgRNA. Middle: Tertiary structure of the *S. pyogenes* sgRNA from 4UN3 (Anders et al., 2014). The spacer (black) is the variable region within the sgRNA that directs spCas9 to specific DNA sequences. The scaffold, made up of the tetraloop (cyan), nexus (orange), and hairpins (lavender), is the structurally conserved element of the sgRNA that functions to bind spCas9 and enable proper conformation prior to DNA binding. Each of these elements of the sgRNA is necessary for spCas9 activity; however, there is considerable flexibility for sequences at the ends of the hairpin and tetraloop elements (red). The design for pGuides is based on insertion of a heterologous sequence within one of the flexible locations (red) to inactivate the sgRNA. G. Structural model of the spCas9-sgRNA-DNA ternary complex (PDB #4UN3). The tetraloop and hairpin 1 protrude from the protein and are exposed on the surface of the riboprotein. The exposure makes both suitable locations for ribozyme insertion, because there are few constraints on the insertion sequence. H. Primary structures of hairpin variant or the tetraloop variant pGuides containing a hammerhead ribozyme (red). Ribozyme cleavage sites are indicated by the red triangle. Sizes of the resulting cleavage products are noted.

To determine optimal locations for a removable inactivating sequence in an sgRNA, we utilized well understood structural and biochemical features of the spCas9-sgRNA ribonucleoprotein (RNP) (Fig 1 E,F). Surfaces of the tetraloop and hairpin 1 structures of the sgRNA are exposed in the RNP structure (Fig 1F), and substantial insertions such as MS2 aptamers have been engineered into these locations without apparent effect on the RNP recognition of target DNA (Konermann et al., 2015). Therefore, we considered these two locations to be quite flexible in terms what RNA sequence could be inserted without interrupting the activity of the RNP. By contrast, sgRNA require both a hairpin 1 structure and tetraloop structure to confer activity in RNP, and truncating sgRNA to remove either hairpin 1 or tetraloop effectively destroyed activity (Briner et al., 2014). Therefore, we designed pGuides to possess a cis-cleaving ribozyme as the inactivation sequence within either the tetraloop site or the hairpin 1 site; this was designed to inactivate the pGuide RNA when it was transcribed, yet allow for multiple modes of inactivating the ribozyme at the DNA level via genome editing (Fig. 1G). To stimulate complete removal of the ribozyme-encoding DNA, ribozymes were flanked by sequences that we call stems, which extend from the hairpin structures, contain targeting sites for other sgRNA, and contain sequences stimulating MMEJ-mediated repair mechanisms in cells (Fig. 1H).

To test the inactivation of sgRNA by cis-cleaving ribozymes, prototype pGuide-encoding DNA were engineered to contain different ribozymes in the hairpin 1 (Fig. S1A). DNA were used as templates for in vitro transcription reactions, and RNA products were resolved on a denaturing gel, demonstrating self-destruction of the RNA molecules into fragments predicted to be non-functional as guide RNAs (Fig. S1B). The activity of these prototype pGuides was tested in cells by targeting them to EGFP with six different spacer sequences (Fig. S1C) and comparing their ability to reduce GFP fluorescence in Rex1::GFPd2 cells relative to sgRNA using the same spacer sequences (Fig. S1D). Each of the prototype pGuides were effectively inactivated by their ribozyme insertions (Fig. S1D); the Hammerhead ribozyme was chosen for further examination because of the combination of its effectiveness and relatively small size.

The Hammerhead ribozyme and the flanking stem structures (Fig. 1H) were inserted into either hairpin 1 or tetraloop site to make pGuides targeting EGFP (Fig. 2A). Analysis of RNA products following in vitro transcription demonstrated that both variants were co-transcriptionally destroyed in vitro (Fig. 2B). Transient transfection of plasmid DNA encoding expression of these pGuides showed that they displayed low levels of sgRNA-like activity relative to an sgRNA control (Fig. 2C). To detect residual activity of these pGuides over a longer duration, they were integrated into the genome of a spCas9- and EGFP-expressing 4T1 cell line using piggyBAC transposition. Activity was assessed by measuring GFP fluorescence with flow cytometry (Fig. 2D) and by quantitative DNA deep sequencing of PCR amplified genomic target sites in the EGFP gene (Fig. S2A-C). Both assays showed that the parental cell line lacking guide RNA expression did not lose GFP fluorescence or gain mutations in the EGFP gene, and the sgRNA-expressing cells lost GFP and gained mutations in more than 90% of cells by the first timepoint after introduction of the sgRNA (10 days) (Fig. 2D). The hairpin variant pGuide was effectively inert during continuous culturing of cells and displayed similarly low levels as the negative control for the 72 day duration of the experiment (Fig. 2D). The tetraloop variant displayed leakiness with a mean 0.36% of cells displaying disrupted EGFP per day of continuous culture of cells (Fig. 2D). Analysis of the pGuide insertions from genomic DNA of long-term cell cultures indicated that the structure of the genomic pGuide DNA insert remained intact and free of mutations within the stem domains (Fig. S2B). DNA sequencing analysis of the target sites in EGFP showed that the top three mutations generated by the sgRNA, including the single base insertion at the −4 position, were also the three most frequent mutation occurring in the tetraloop variant by day 72 (Fig. S2C). Although the DNA encoding the pGuide did not indicate a conversion to an active state had occurred, the low level of leaky activity in the tetraloop variant was consistent with normal sgRNA activity indicating that pGuide mediated mutagenesis was spCas9-induced. We suggest the low level of leakiness is caused by either infrequent failure of the ribozyme to cleave the sgRNA before RNP complex is formed or by residual RNA fragments having a weak ability to form an active RNP complex in cells. Taken together, these data demonstrate that the ribozyme insertion effectively disrupts the guide RNA and prevents its spCas9-related activity in cells.

**Figure 2:**
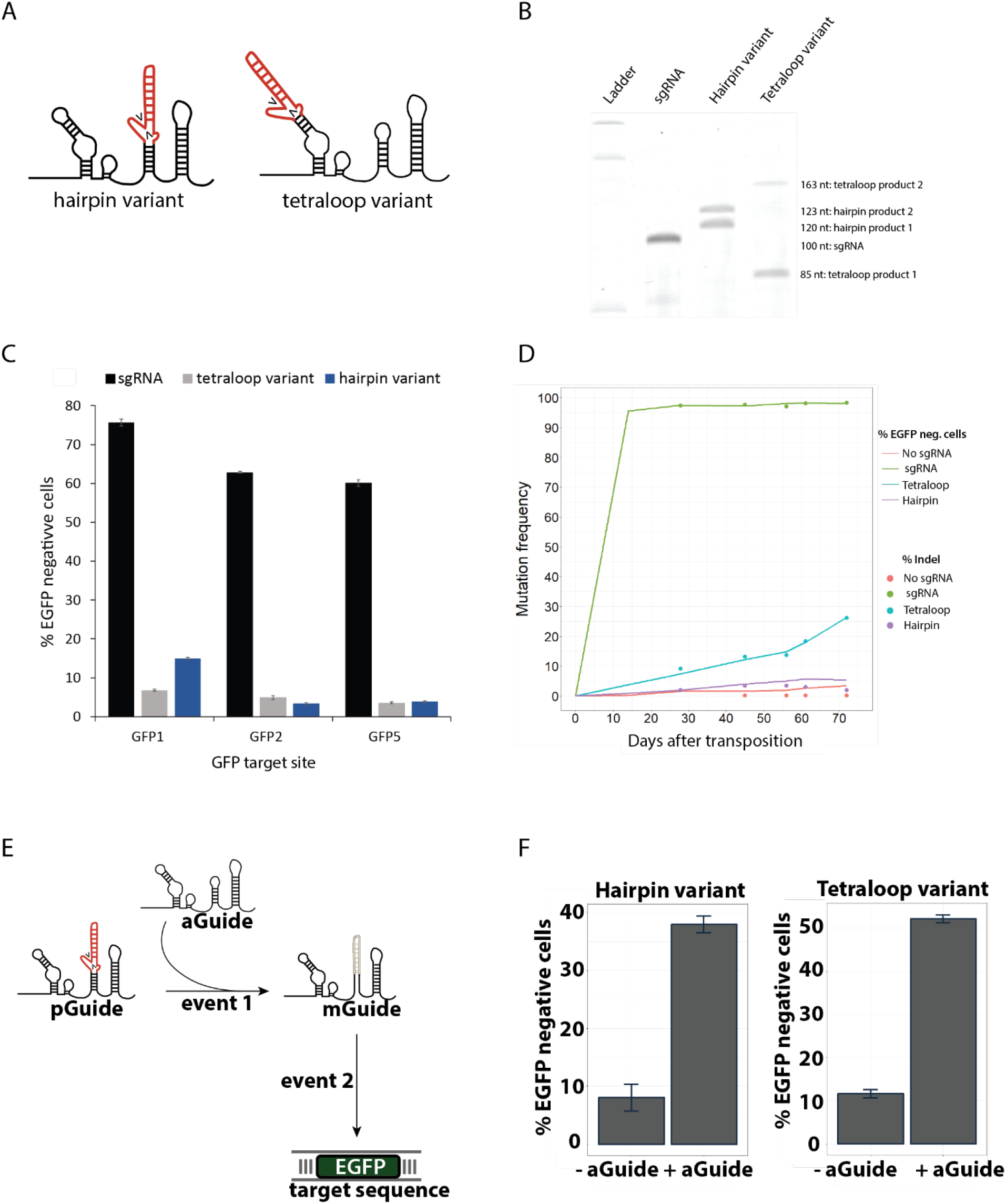
Self-destructing pGuides provide latent Cas9 targeting activity upon conversion by an sgRNA. A. Schema depicting two pGuide designs possessing ribozyme encoding sequence (red) at either the end of the first hairpin (hairpin variant) or at the end of the tetraloop domain (tetraloop variant). Arrowheads depict cleavage of the pGuide RNA by the ribozyme after transcription. B. sgRNA and pGuide DNA were subjected to *in vitro* transcription and sizes of RNA products were analyzed via EtBr staining following electrophoreses through a TBE-urea denaturing gel. Intact hairpin variant pGuide RNA (243nt) and intact tetraloop variant pGuide RNA (248nt) were not detected in the gel. Ribozyme cleavage products of pGuides (sizes indicated on right) and intact sgRNA (100nt) were present. Note, product 1 refers to the RNA fragment from the 5’ end of the ribozyme cleavage, and product 2 refers to the RNA fragment from the 3’ end of the ribozyme cleavage. See Fig. S1A,B for comparison of other ribozymes. C. Cellular inactivation of pGuide by the ribozyme was assessed by comparing the ability of pGuides and sgRNA targeting a genomic EGFP gene. 4T1::EGFP cells harboring a single genomic insertion of an EGFP expression gene were transiently transfected with a spCas9 expression plasmid and either a sgRNA (black) or a pGuide (gray; tetraloop variant, blue: hairpin variant) expression plasmid. EGFP disruption at three different sites (GFP1, GFP2, GFP5) was tested by using pGuides or sgRNA targeted to each of the corresponding DNA sequences (Fig. S1C, Table S1) in separate transfections. Five days after transfection, cells were analyzed by flow cytometry to determine percentage of cell lacking GFP fluorescence. Data are displayed as the mean +/−standard deviation of n = 3 biological replicates. D. Long-term latency of pGuide activity was examined in continuous cell culture of spCas9-expressing 4T1::EGFP cells that were subjected piggyBac transposition of negative control (no sgRNA), sgRNA, or pGuide (Tetraloop or Hairpin) expression DNA. The pGuide and sgRNA targeted the same sequence in EGFP, and activity was assessed by measuring GFP fluorescence by flow cytometry (line) and by deep sequencing of the target site in genomic DNA isolated from cells (points). See Fig. S2A,C for PCR and deep sequencing analysis of the pGuide genes. Data is representative from a starting polyclonal population of ~1,000,000 cells during 72 days of continuous cell culture. E. Schema illustrating the conversion of a pGuide into an mGuide that targets EGFP. F. Conversion of latent pGuide to active mGuide state in long-term cultures of cells in Fig. 2D as illustrated in (E). After 28 days of culture, cells harboring each pGuide variant (Hairpin and Tetraloop) were transiently transfected with an aGuide expression plasmid (+aGuide) or empty expression plasmid (-aGuide) control. Loss of GFP fluorescence was measured by flow cytometry five days after transfection. Data are displayed as mean +/−range of n = 2 biological replicates.

### Conversion of pGuides to mGuides by genome editing

In addition to remaining in an inactive state, a primary requirement for effective an pGuide is conversion into an mGuide, which direct spCas9 activity to its cognate target sequence. Because it has a singular function for converting a pGuide to an active state, hereafter we use the term “activating-Guide” (aGuide) to refer to an sgRNA that targets a pGuide for conversion (Fig. 1D, 2E). By this definition, aGuides do not target other genomic sites. To test the feasibility of pGuide conversion, we transfected aGuide-expressing plasmids into the spCas9- and pGuide-expressing 4T1 cell lines after they had been continuously cultured for 28 days (Fig. 2E). Conversion to an active mGuide state was assessed by loss of GFP expression five days after transfection (Fig. 2E, F). Whereas control transfections without aGuide displayed low levels of GFP-negative cells, addition of the aGuide caused a loss of GFP fluorescence in 40% of cells harboring the hairpin variant pGuide and 48% of cells harboring the tetraloop variant pGuide (Fig. 2F). These results suggest after about a month of cell culture, that the aGuide successfully stimulated the conversion of genomically integrated pGuides to an active mGuide state, which targeted spCas9 activity to the genomic site corresponding to the spacer sequence in the mGuide.

To develop a pGuide system of preprogrammed sequential genome edits, it was important to elucidate the capabilities and limitations of pGuides and how they function as sgRNA-like molecules. Therefore, the efficiency, kinetics, and mutational outcomes of genome editing stimulated by pGuides were compared to those stimulated by sgRNA. Experiments were conducted primarily in Rex1::GFPd2 mouse embryonic stem cells, because the destabilized EGFP gene integrated into the Rex1 locus produces GFP molecules with a half life of ~2 hours (Wray et al., 2011), which improves analysis of kinetics of loss of GFP fluorescence following spCas9-mediated mutation of EGFP. The pGuides targeting EGFP (as described in Fig. 2) were integrated into the genome of Rex1::GFPd2 cells via transposition, and experiments were conducted by transiently transfecting an spCas9-expressing plasmid with or without an aGuide plasmid. As a reference, pGuide effects were compared to the Rex1::GFPd2 cells transiently transfected with the spCas9-expressing plasmid and an sgRNA plasmid. Flow cytometry analysis of GFP levels after 48 hours showed that sgRNA disrupted GFP in 28% of cells and addition of an aGuide disrupted GFP in 4% and 5% of hairpin and tetraloop pGuide-containing cells, respectively (Fig. 3A). For all conditions, frequencies of GFP disruption increased over time, and sgRNA mediated GFP disruption reached a final frequency of 70% of cells while tetraloop and hairpin pGuide cell lines transfected with aGuides reached 35% and ~4%, respectively (Fig. 3A). Thus, conversion of the genomically integrated pGuides resulted in about half the overall activity of a plasmid-expressed sgRNA by 120 hours.

**Figure 3:**
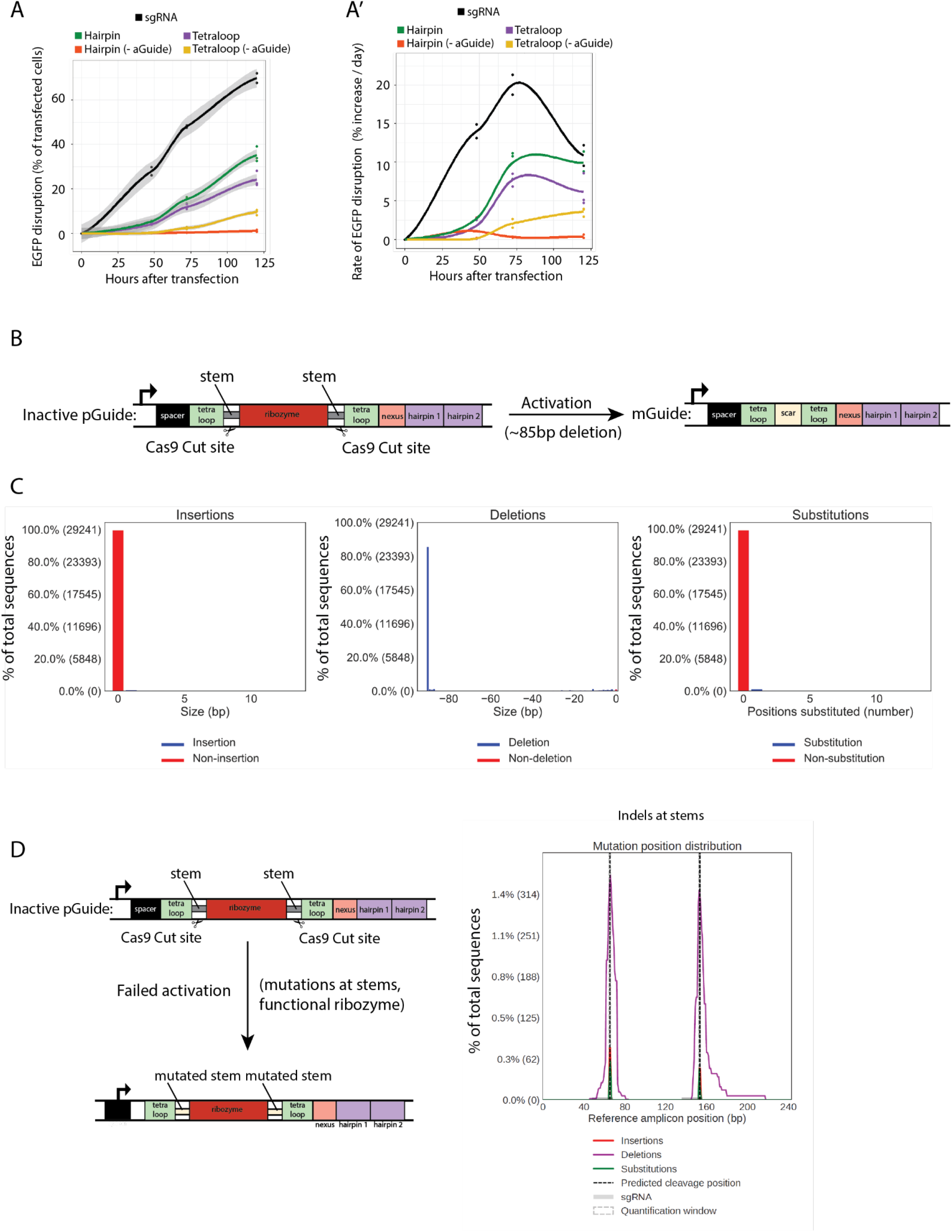
Conversion of pGuide to an active mGuide state by removal of ribozyme encoding DNA. A, A’. The overall experimental approach is illustrated in Fig. 2E. The activity of mGuides was measured by loss of EGFP expression from Rex1::GFPd2 cells harboring genomically integrated pGuide DNA at indicated times after transient transfection of aGuide expression plasmid. EGFP disruption was assessed by loss of GFP fluorescence detected by flow cytometry. A: EGFP disruption values over 125 hours for tetraloop pGuides +/−aGuide, hairpin 1 pGuides +/−aGuides, and the positive control sgRNA targeting EGFP. A’: Rate of EGFP disruption as a function of time for the data in (A). Data are displayed as dots representing n = 3 biological replicates, lines being the average of all three dots for each condition, and gray shading representing the standard deviation. B. Schematic depicting the process of ribozyme excision via genome editing. Identical target cut sites flank the ribozyme within sequences called “stems.” The example shown depicts excision of the ribozyme from the tetraloop variant; the same design was applied to the hairpin variant. Excision of the ribozyme via spCas9 cutting at each stem and repair by end joining results in a deletion of 85bp. Note that the stems are designed to favor this large excision to be the predominant repair event in cells. C. Stem sequences stimulate deletion of entire ribozyme from a genomically integrated pGuide. A tetraloop variant pGuide (diagrammed in A) was integrated into the genome of mESC::Rex1 cells. These cells were transiently transfected with plasmids expressing spCas9 and an aGuide, and genomic DNA was isolated 40 hours after transfection. The pGuide/mGuide was amplified via PCR and subjected to deep sequencing. The predominant mutation detected was an 85 bp deletion (middle graph, blue bars). Data represent distribution of 29,241 DNA sequencing reads from one of two replicates. D. Non-converting indel mutations within the stems sequences occur via indel mutation at each stem sequence without deletion between the two stems (Left). Deep sequencing showed that double indels was a rare event, occurring in less than 1.5% of the 29,241 sequencing reads. Data is representative from one of two replicates.

The kinetics of gene editing was examined by comparing EGFP disruption for each day during the 120 hr duration of the experiment. As expected, the sgRNA generated a relatively linear increase in EGFP disruption after transfection at an average rate of 24% editing per 24 hours until it peaked at 72 hours (Fig. 3A’). After the peak, rates of further EGFP disruption by the sgRNA decreased to 11% per 24 hours as the number of unmutated substrates diminished in the cell population (Fig. 3A’). By contrast, aGuides stimulated a different kinetic profile in pGuide-expressing cells. Compared to sgRNA, pGuide conditions exhibited substantially slower EGFP disruption rates of 2.7% and 2.1% per 24 hours for tetraloop and hairpin pGuides respectively during the first 48 hours (Fig 3A’). Interestingly, from 48 to 72 hours, the rates from pGuides increased to 15% and 11% per day. Notably, the increase in the rate of GFP disruption from 48 to 72 hours for pGuides, which were similar to the 14% per 24 hours rate mediated by sgRNA for hours 0 to 48. (Fig 3A’). Finally, the rate of GFP disruption for pGuides slightly increased from 72 to 120 hours, with tetraloop pGuides editing at 17% per 24 hours and hairpin variants at 12% per 24 hours. These data indicate that, as expected, pGuides stimulate genome editing with complicated kinetics that include a delay period during which conversion to an mGuide occurs, followed by an active period with kinetics similar to an sgRNA with an extended period of peak editing rates. This is notably distinct from other ways of prolonging the duration of sgRNA activity, such as by incorporating mismatches to their spacer sequence (Chan et al., 2019), which display a standard sgRNA profile albeit at slower rates.

Conversion of pGuides into an active mGuide state requires inactivation of the cis-cleaving ribozyme. With the inclusion spCas9-target sites on both sides of the ribozyme sequence in stem regions, the conversion is designed to occur by deletion of the entire ribozyme nested between the stem regions (Fig. 3B). To examine the mutagenesis profile associated with pGuide to mGuide conversion, we harvested genomic DNA from the entire cell population for each condition in Fig. 3A-A’’ at the 96 hours timepoint, PCR amplified the pGuide insertions, and analyzed them by deep sequencing. Expression of aGuides stimulated primarily deletions between the two spCas9 cut sites in the stems (Fig. 3C). By contrast, indel mutations within a stem would fail to inactivate the ribozyme and fail to stimulate conversion; however, such non-converting edits occurred infrequently (Fig. 3D). This data suggests that ribozyme excision via aGuides is responsible for pGuide to mGuide conversion.

### Episomal pGuide systems

To construct cascades of multiple pGuides, the spacer sequence from one pGuide must target both stems in a downstream pGuide. Empirically testing those sequences is necessary for constructing cascades of pGuides, because the targeting efficiency and repair outcomes of spCas9 on target sequences cannot be predicted perfectly. To facilitate testing of pGuide sequences, we sought to determine if pGuides could be converted to mGuides encoded by episomal plasmid DNA (Fig. 4A), and not require integration into the genome. In a single transient transfection, the plasmid DNA for the tetraloop variant pGuide from Figure 2, its aGuide, and spCas9 expression were introduced into Rex1::GFPd2 cells. The frequency of GFP disruption via conversion of the pGuide was similar for the episomal pGuide plasmid relative to the genomic pGuide (Fig. 4B). Although integration of pGuides into the genome enables long term latency (Fig 2D), most experiments in the remainder of this study use pGuides encoded on plasmid DNA because they obviate the requirement to engineer cell lines for experiments. In addition, episomal pGuide cascades provide potential in applications that are sensitive to insertion of foreign DNA into a cell’s genome.

**Figure 4:**
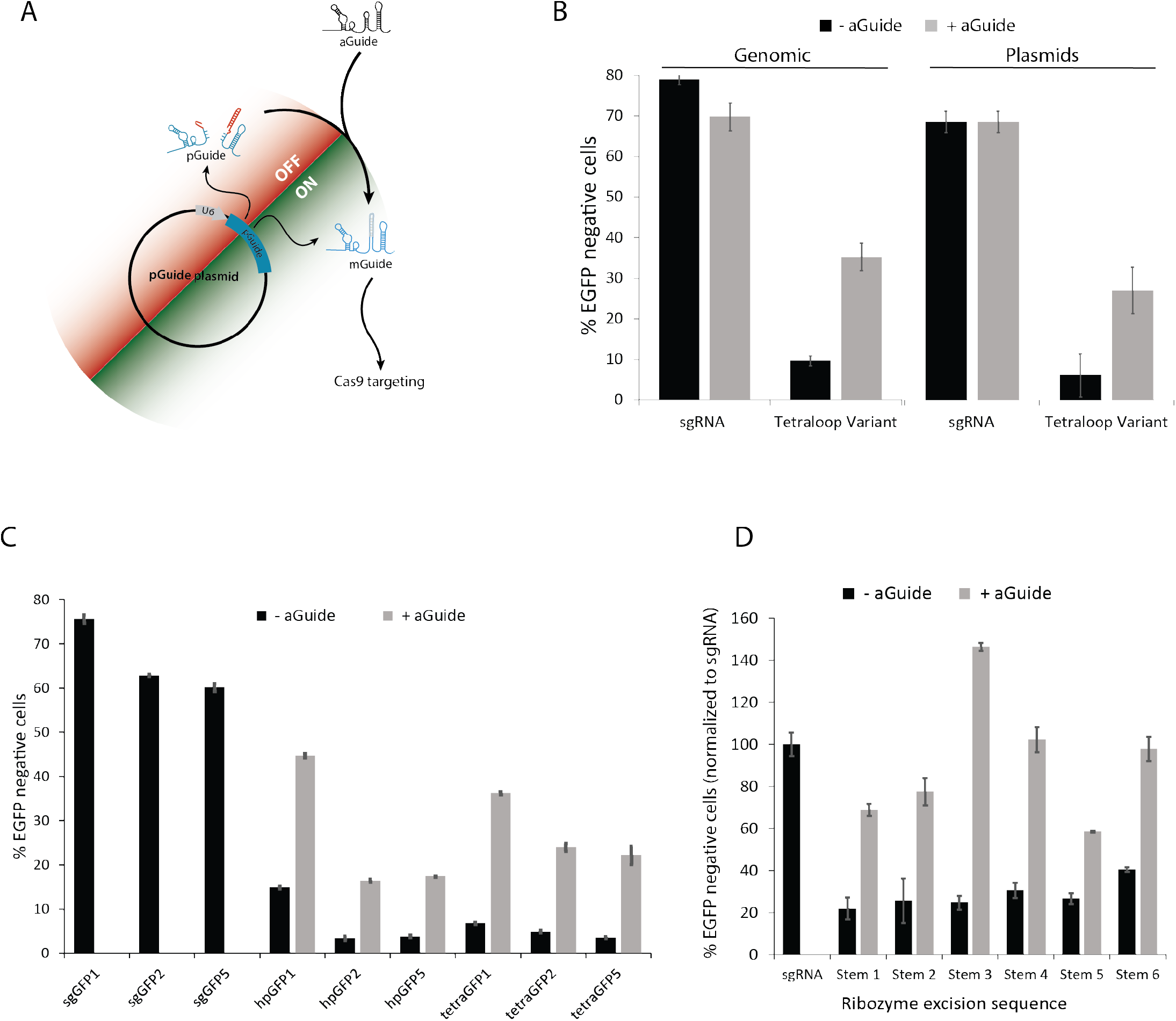
Utility of plasmid based pGuides. A. Schema highlighting steps in the conversion of an episomal pGuide to an mGuide via an episomal aGuide. Elements of the conversion are provided via transient transfection of plasmid DNA, which do not require genomic integration for their activity. Multiple copies of a pGuide plasmid causes cells to express both latent pGuide (OFF) and activated mGuide (ON) RNA following activity of the aGuide. The presence of the mGuide RNA targets spCas9 to the genome. B. The relative effectiveness of a genomically integrated pGuide (Genomic) and an episomal pGuide (Plasmids) was assessed in Rex1::GFPd2 cells by loss of GFP fluorescence following transient transfection of plasmid DNA. As a positive control, an sgRNA targeting the same sequence in EGFP was transfected without pGuides. Exclusion of the aGuide plasmid from transfections was used to assess activity that was dependent upon conversion of the pGuide. Data are displayed as the mean +/− standard deviation of n = 3 biological replicates. C. The effect of the mGuide target site on its genome editing activity was assessed for three different target site sequences (GFP1, GFP2, GFP5; see also Fig. S1C and Table S1). spCas9-, pGuide-, and aGuide-expression plasmids were transiently transfected into Rex1::GFPd2 cells, and GFP disruption was measured three days after transfection. The sgRNA and –aGuide controls are as described in (B). Data are displayed as the mean +/−range of n = 2 biological replicates. D. The effects of elements of the stem sequences on conversion of pGuides was assessed for six different stem sequences containing different spCas9 target sites (see Fig. 1G, H and Table S1 for nucleotide sequences). Conversion was measured by targeted each pGuide to EGFP and assessing disruption frequency as in B and C. Relative EGFP disruption frequencies were calculated by dividing EGF disruption from the pGuide conditions by EGFP disruption frequencies mediated by the sgRNA positive control. Data are displayed as mean +/−the range for n = 2 biological replicates.

The importance of the location of the ribozyme in episomal pGuides, either in hairpin 1 or tetraloop location, was examined by testing conversion of different episomal pGuide plasmids targeting three different sites in EGFP. Within 72 hours, each of the pGuide plasmids was converted to an active state, resulting in disruption of the genomic EGFP gene in Rex1::GFPd2 cells (Fig. 4C). Conditions lacking aGuides displayed low frequencies of GFP disruption. The level of GFP disruption correlated with the spacer sequence, as the best performing spacer for sgRNA (GFP1) was also superior for the hairpin and tetraloop variant pGuides, and the least effective spacers for sgRNA (GFP2, GFP5) displayed lowest levels in pGuides (Fig. 4C).

To test whether the DNA sequence in the stem significantly affects pGuide conversion, six pGuides with distinct stems were evaluated for aGuide-mediated conversion to an active, GFP-disrupting, mGuide state (Fig. 4D, S3A). Each pGuide possessed the same spacer sequence (GFP1), but they differed by the cut site sequences embedded in the stems and the orientation of the cut site with respect to which strand possessed the NGG PAM sequence (Table S1). A significant range of GFP-disruption was exhibited by the different stems; the least effective stem sequence disrupted GFP at only a 59% level relative to the sgRNA control, and the most effect stem disrupted GFP at a 146% level, better than the sgRNA control (Fig. 4D, Fig. S3A). These data suggest the importance of nucleotide composition of stem sequences within the pGuide for determining their capacity for conversion to an mGuide state.

### Constructing and optimizing multi-pGuide cascades for more than two sequential gene edits

A critical characteristic of the pGuide design is that the conversion of one pGuide to an mGuide can be used to stimulate the conversion of another pGuide in a sequential cascade of events (Fig 1). To test the capabilities of pGuides to function in such a modular and sequential manner, several prototype cascades pGuides were generated using the set of validated pGuide stem variants (Fig. 4D, S3A, Table S1). First, linear cascades consisting of one- or two - pGuides were generated whereby one pGuide module activates the next in a daisy chain arrangement ending with an mGuide targeting disruption of a genomic EGFP gene (Fig. 5A, S4A). GFP disruption frequencies were measured for a one-pGuide cascade and two arrangements of two-pGuide cascades 96 hours after transfection and were compared to frequencies achieved by an sgRNA targeting the same site as the final mGuide (Fig. 5B, S3B). Interestingly, one of the two pGuide cascades significantly disrupted GFP (Cascade 1), and the latent activity in the conditions lacking aGuides was significantly less than the successful two-pGuide cascade (Fig. 5B, S3B). Interestingly, the one-pGuide cascade and was not significantly different from the more effective two-pGuide cascade at the end of the 96 hr course of this experiment.

**Figure 5:**
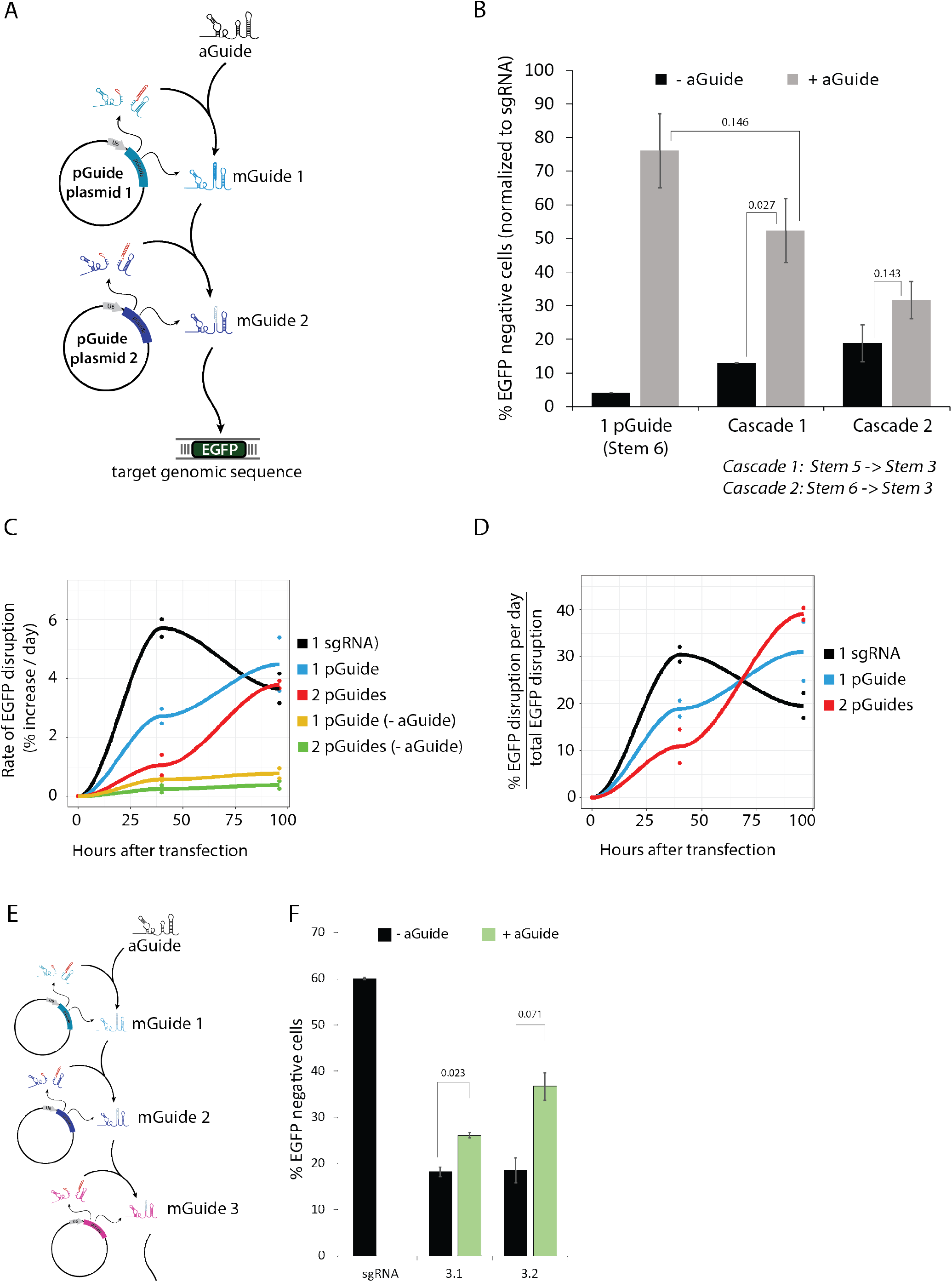
Sequential spCas9 activities via pGuide cascades display stepwise, delayed kinetics. A. Schema of a linear three-step, two-pGuide cascade that targets disruption of a genomic EGFP via these three spCas9-mediated activities: 1) conversion of pGuide 1 by aGuide, 2) conversion of pGuide 2 by mGuide 1, and 3) disruption of EGFP by mGuide 2. B. Plasmid-based, two-pGuide linear cascades were transiently transfected into Rex1::GFPd2 cells, and disruption of GFP fluorescence was measured by flow cytometry. For comparison, a single pGuide (as in Fig 3) targeting GFP was included in the experiment. The two linear cascades used a different stem sequence in pGuide plasmid 1 and displayed different levels of EGFP disruption. EGFP disruption frequencies for all pGuide conditions were divided by EGFP disruption frequencies mediated by sgRNA positive control to produce normalized values (see also Fig. S4B). Data are displayed as mean +/−standard deviation of n = 3 biological replicates. p-values are located above lines drawn between two conditions and were calculated using Welch’s two sided t-test. C. Kinetics of EGFP disruption are plotted for the one-pGuide and two-pGuide cascade (Cascade 1) from (B). The rate of editing per day was determined through calculating the total absolute frequency of editing per 24-hour period (e.g. first 24 hours, second 24 hours, etc). 1 sgRNA represents a single genome editing event, 1 pGuide represents 2 genome editing events, and 2 pGuides represents three genome editing events. Dots represent two biological replicates and the line is an average of the two replicates for each condition. D. Rate of editing from (C) plotted as a percent of max for each condition by calculating the total percent editing for every 24-hour segment and dividing it by the final editing frequency (e.g. % editing for 0-24hours / total editing) in order to normalize each condition to 100%. E. Schema illustrating a plasmid-based, three-pGuide linear cascade executing four sequential spCas9-mediated events. The final mGuide targets EGFP. F. Completion of the linear cascade in Rex1::GFPd2 cells was assessed by EGFP loss by flow cytometry as described above, except a Cas9Cdt1 expression plasmid was used in place of spCas9. Data are displayed as mean +/−s.d. for n = 3 biological replicates. p-values were calculated using Welch’s two sided t-test.

Given that the two-pGuide cascades require more genome editing events than a one-pGuide cascade or an sgRNA, we anticipated a delay in the disruption of GFP from cascades relative to the sgRNA. Therefore, the kinetics of the effective two-pGuide cascade were examined by measuring GFP disruption at an earlier, 48 hr timepoint. Whereas the rate of GFP disruption peaked for sgRNA at 48 hrs, the one- and two-pGuide cascades did not display a decrease in the rate of GFP disruption after 48 hrs (Fig. 5C). Consistent with the delay observed from genomically integrated pGuides (Fig 3A, A’), the episomal one-pGuide cascades in this experiment were delayed relative to sgRNA; however the delay for the episomal pGuides was not as long, as the one-pGuide cascade displayed significant GFP disruption by 48 hours (Fig. 5C). The two-pGuide cascade displayed low levels of GFP disruption at 48 hours, and its rate of disruption increased rapidly from 48 to 96 hours (Fig. 5C). When normalized for overall GFP disruption frequency at 96hr for each cascade, substantial differences in editing rates are apparent; the sgRNA displayed the highest rate from 0-48 hrs and the two-pGuide cascade was highest from 48-96 hrs (Fig 5D). These analyses are consistent with the genome editing events encoding by pGuide cascades proceeding in a ratchet-like, stepwise manner.

Initial attempts to construct three-pGuide cascades, we initially used unmodified spCas9 and episomal pGuide plasmids (Fig. 5E, S4). Only one of the cascades yielded a small, but statistically significantly increased GFP disruption relative to the no aGuide control (Fig. S4). Considering that the addition of genome editing steps in longer cascades of pGuides resulted in longer delays in in GFP disruption, it was possible that the three-pGuide cascade was unable to progress to completion with the 96 hour timeframe and/or the number of guide RNA molecules competing for spCas9 reduces the efficiency genome editing. The stable binding of spCas9 to its guide RNA and the stable binding of the RNP to DNA after generating a DNA break are two possible rate-limiting steps in the progression through a cascade of pGuides (Clarke et al., 2018; Sternberg et al., 2014). Notably, previous studies have reduced the half-life of spCas9 to increase the efficiency of precise excision (Tu et al., 2017; Yang et al., 2018).

Therefore, we fused the Fucci degrons from Geminin and Cdt1 to the carboxy terminus of spCas9 to generate destabilized spCas9 proteins termed Cas9Gem and Cas9Cdt1 (Fig. S3C)(Sakaue-Sawano et al., 2008). The effects of destabilizing spCas9 was tested on pGuide cascades. Cas9Cdt1 displayed 150% the activity of unmodified spCas9 (WTCas9) for GFP disruption from a one-pGuide cascade (Fig. S3D). By contrast, Cas9Gem displayed 50% the activity of unmodified spCas9. The addition of the degron sequences to spCas9 did not appear to affect the types of mutations that accumulated in the EGFP site targeted by the mGuide (Fig. S3E). When used with the episomal three-pGuide cascade, Cas9Cdt1 increased the disruption of GFP relative to unmodified spCas9 for both Cascade 3.1 and Cascade 3.2 (Fig. 5E, F). Together, these results demonstrate capability of a linear cascade of four genome editing events encoded entirely by episomal plasmid DNA to progress to completion and target the genome in mammalian cells.

### Ramified pGuide cascades for parallel and sequential spCas9 activities

Although daisy-chain style pGuide cascades (Fig. 5A, 5E) progress in a linear manner with one pGuide encoding the conversion of the next pGuide, non-linear configurations of pGuide cascades are also possible and would enable substantial additional functionality. Branch points that ramify pGuide cascades can be introduced by encoding the conversion of multiple pGuides to occur via the same target sites in stem domains. We designed a ramified cascade of pGuide plasmids to target two different genomic sites in the host cell, which would result in the sequential acquisition of two unrelated phenotypes: ouabain resistance and RFP expression (Fig. 6A). The first genomic site was the ATP1A1 gene, which confers sensitivity to the drug ouabain, and can be edited to confer ouabain resistance via a previously characterized gene editing procedure (Fig. S5A) (Agudelo et al., 2017). Targeting of pGuides to ATP1A1 stimulated resistance to ouabain as measured via calcein AM staining when cells were co-transfected with an aGuide (Fig. S5B, C). The second genomic site uses an out of frame RFP coding sequence to detect indel mutations upstream of RFP that result in the coding sequence to be placed in frame, a system described previously as the traffic light reporter (TLR) (Fig. S5D) (Chu et al., 2015). Three sgRNA were tested for their ability to restore the RFP reading frame (Fig. S5D, E), and the spacer sequence from the most effective sgRNA was used to target the pGuide to the TLR genomic site (Fig. S5F, G). The prototype ramified cascade places the ATP1A1 edit one step earlier than the TLR edit, but in a parallel branch (Fig. 6A). The conversion of the pGuide targeting ATP1A1 to an mGuide is not required for conversion of the pGuide targeting TLR (Fig. 6A).

**Figure 6:**
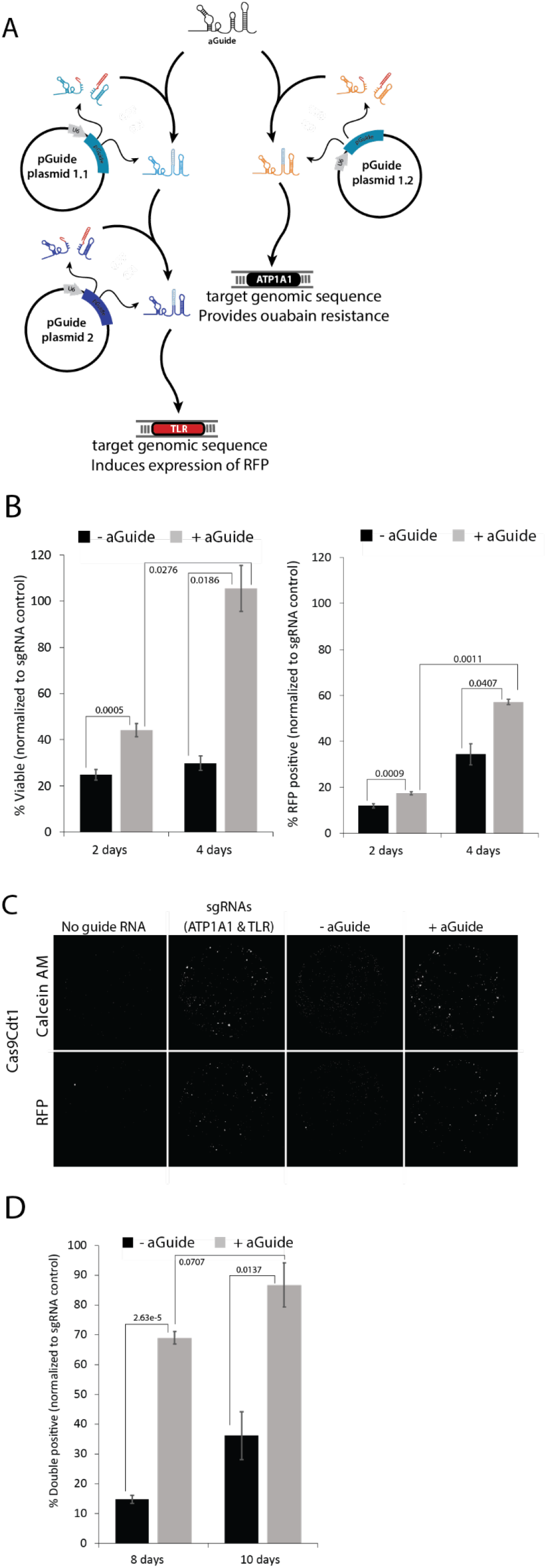
Programming temporal ordering of spCas9 activities via a genome-independent, plasmid-based cascades of pGuides. A. Schematic of a ramified five-step pGuide cascade that targets the genome at two sites (TLR and ATP1A1) through execution of these five spCas9-mediated activieis: (left side) 1) activation of pGuide 1.1 by aGuide, 2) activation of pGuide 2 by mGuide 1.1, 3) editing of TLR gene by mGuide 2 to induce RFP expression, and (right side) 4) activation of pGuide 1.2 by aGuide, and 5) editing of ATP1A1 gene by mGuide 1.2 to provide ouabain resistance. See Fig S5B-C for pGuide editing the ATP1A1 gene and Fig. S65F-G for pGuide editing the TLR gene. B. Left: Plasmid DNA constituting the ramified cascade and Cas9Cdt1 shown in (A) were transiently transfected into HEK293 cells harboring a genomic copy of TLR. Progression to the two genomic endpoints was assessed by resistance to 0.5uM ouabain for the ATP1A1 gene and RFP fluorescence for the TLR gene. Resistance to ouabain was determined by viable cell staining with Calcein AM after six days of selection in ouabain-containing media. Right: Percent viability was determined by dividing the number Calcein AM positive cells after six days of ouabain treatment by the number of starting cells plated at two days or four days after transfection. The percent RFP positive was determined by measuring individual cell fluorescence with a Celigo instrument within an hour of plating cells (two and four days). Data are normalized to levels of viable cells and RFP positive cells generated by transfection of sgRNA targeting ATP1A1 and TLR, respectively, from concurrent transfections. See Fig. S6 for data prior to normalization. Data represent means +/−standard deviation of n = 3 biological replicates. p-values are located above or below the lines drawn between two conditions and were calculated using Welch’s two sided t-test. C. Representative images of cells that underwent six days of ouabain treatment initiated two days after transfection of plasmid DNA. Fluorescent images of entire well from a 96 well plate using a Celigo instrument. Images are representative of three biological replicates. D. Percentage of cells that were resistant to ouabain starting at two days and four days and have progressed to become RFP positive by the end of ouabain treatment at eight days and ten days after transfection. Viability and RFP positivity were measured as described in B. Data are normalized to levels of viable cells and RFP positive cells generated by transfection of sgRNA targeting ATP1A1 and TLR, respectively, from concurrent transfections. See Fig. S6 for data prior to normalization. p-values were calculated using Welch’s two sided t-test.

To test completion of the cascade at the two end points, a single aGuide plasmid was co-transfected with three pGuide plasmids and a Cas9Cdt1 expression plasmid (Fig. 6A). Progression to the endpoints in the ramified cascade was assessed by comparison to control cells transfected with sgRNA directly targeting TLR or ATP1A1 (Fig. 6B, C). Resistance to ouabain developed more slowly from the pGuide cascade compared to the sgRNA; the pGuide cascade required 96 hours to reach levels the sgRNA had reached after 48 hours (Fig. 6B), which is consistent with the lag observed from pGuide relative to sgRNA in linear cascades (Fig. 5C). The appearance of RFP fluorescence in the population was slower than ouabain resistance in cells transfected with the cascade (Fig. 6B,C) and slower than RFP fluorescence in cells transfected with sgRNA (Fig. S6A). The slower accumulation of the TLR edit via the cascade is consistent with the requirement for two pGuide maturation steps compared to one (for ATP1A1) or none (for sgRNA control). In cells that were selected for ouabain resistance for the final six days, a high frequency of TLR edits was observed (Fig. 6C,D). Importantly, >70% of cells that were ouabain resistant were also RFP positive by the end of the timecourse. As time progressed, the ouabain resistant cell population accumulated RFP positivity, supporting the sequential activity of the pGuide cascade. These observations are consistent with a stepwise progression through individual modules of the cascade whereby the timing of endpoint event can be controlled by the number of pGuides placed between the aGuide and the endpoint.

## DISCUSSION

The scale and ease of multiplexing spCas9 activities has provided transformative-level capabilities to synthetic biology and genetic engineering in mammalian cells. However, capabilities have been limited by a lack of methods that can control either when or under what conditions individual multiplexed events occur relative to one another. In this study, the pGuide has been developed to be modular tool for programming interdependent, sequential, and possibly temporal controls over multiple spCas9-mediated events in individual cells. Prototype pGuides remained inactive for over a month in mammalian cells until they were triggered to undergo a conversion to an active state by an spCas9-mediated gene editing event. Upon conversion to an active mGuide state, they appeared to function similarly to sgRNA in directing spCas9 activity. Moreover, by designing the spacer sequence of one pGuide to target another pGuide for conversion to an mGuide, proof-of-principle cascades were generated encoding five genome editing events with either a linear daisy-chain or a ramified architecture. The functionality of these cascades, encoded on episomal plasmid DNAs, illustrate the potential for pGuides cascades to be assembled for a wide range of applications requiring preprogrammed ordering of multiple spCas9-mediated activities.

The ability to preprogram the sequential activation of spCas9 at multiple genomic sites introduces a new tool for biological research and genetic engineering. The passage of time is an integral component of human development and disease progression; however, the molecular and genetic methods used to examine biological systems are often inadequate at addressing time as an important dimension. By enabling progression through pre-programmed cascades of pGuides, the technology presented in this study can enable scientists to address time in new ways. For example, the order by which oncogenic mutations arise in individual cells may affect important characteristics of the resulting cancer; daisy-chain cascades of pGuides could be used to generate mutations in various predetermined sequences of genome editing events to elucidate the effects order. Current lineage tracing using spCas9 takes advantage of the ability to record information about the cell’s origin by making changes to its genome (Chan et al., 2019; Kalhor et al., 2018; Kalhor et al., 2017; McKenna et al., 2016). By offering control over when individual recording events occur relative to one another, a cascade of pGuides would considerably expand the power of current systems to record longer cell lineages through more generations of cells. Other systems have demonstrated the ability to record in a cell’s genome the occurrence of activity from a signal transduction pathway (Frieda et al., 2017; Perli et al., 2016). Cascades of pGuide could be used to provide a timestamp, indicating when the signaling pathway was active relative to progression through a cascade of pGuides.

Although not explored in this study, it is worthwhile noting that spCas9 has been adapted for uses beyond generating a double stranded break for genome editing and that fusing spCas9 with other proteins has been effective at targeting genomic sites with various activities (transcriptional activation, transcriptional repression, histone modifications, fluorescent probes) (Chavez et al., 2015; Hilton et al., 2015; Knight et al., 2015; Larson et al., 2013). Transcriptional activation of target genes has been accomplished by fusing a variety of activation domains not only to nuclease-dead spCas9 (dCas9) (Chavez et al., 2015) but also to nuclease-active spCas9 with Cas9-VPR (Kiani et al., 2015). Combining Cas9-VPR with a cascade of pGuide could enable sequential control of gene expression as the VPR transactivator would be recruited to new genomic sites as pGuides are converted to mGuides in the cascade. As such, it may be possible to use pGuide cascades with Cas9-VPR for sequential epigenetic programming.

Retaining the flexibility of targeting genomic sequences is a critical feature of pGuides and one that distinguishes it from other ratcheting systems, such as self homing guide RNAs (Kalhor et al., 2017; Perli et al., 2016) and DOMINO (Farzadfard et al., 2019). Ideally, any DNA sequence could be used in a stem and be compatible with both conversion to an mGuide state and targeting the cell’s genome for editing, and bespoke cascades of pGuide could be simply engineered for each application. Indeed, the analysis of six different stem sequences validated that they each caused both the inactivation of the pGuide prior to aGuide expression and the conversion to an mGuide upon aGuide expression. However, the efficiency of conversion of different pGuides varied and appeared to depend upon both the spacer sequence and the stem sequence. Nucleotide composition is known to greatly influence DNA repair outcomes at spCas9 generated DSBs (Allen et al., 2018; Chakrabarti et al., 2019; Leenay et al., 2019; Shen et al., 2018; van Overbeek et al., 2016). It is possible that some stem sequences are less amenable to the desired MMEJ repair event if cryptic microhomologies occur because of the nucleotide composition of the spacer region or between the target site in the stem and the ribozyme DNA sequence. It is likely that some combinations of spacer and stem sequences will be incompatible with pGuide conversion. Investigation of the nucleotide parameters influencing pGuide to mGuide conversion is an important next step for reliable engineering of pGuide cascades.

The ability to ramify pGuide cascades provides additional functionality and potential safeguards for overcoming low efficiency cascades. One obvious benefit from ramified cascades is that it allows for more than one genome editing activity to be triggered at the same preprogrammed time. This allows for more sophisticated programs of events to be encoded by pGuide cascades. During testing of daisy-chain and ramified proof of principle cascades, some combinations of pGuides were less effective than others, introducing the possibility that generating bespoke cascades for every application could be limited based on compatibility of nucleotide sequences in stems and spacers. Because inefficiencies at one step would affect completion of all downstream steps in a cascade, a ramified cascade strategy may be needed to retain the critical factor of flexibility in targeting the genome with mGuides. In this case, a single base system of a daisy-chain pGuide cascade would be required to progress to completion without interfacing with the host cell’s genome. In order to target the cell’s genome, ramifying pGuides would be generated for each new application to be converted by the base system cascade at the desired time. Inefficiencies associated with ramifying pGuides would not affect progression through the cascade, which would occur independently of the activity of ramifying mGuides.

In comparison to transcription factor-based approaches to engineering synthetic programs of cellular activities, spCas9-centered approaches offer several advantages. Because these systems use only a single protein (spCas9) to mediate each step in a cascade, the level of sophistication is substantially reduced for engineering a customizable genetic program for mammalian cells. Since each step in a pGuide cascade is encoded by less than 200bp of DNA, cascades can be easily delivered to cells that express spCas9. Although optimization of nucleotide sequences will be required for reliable generation of pGuide cascades, that process is amenable to higher throughput and quicker turnover of iterative optimization steps than the process of engineering protein characteristics. Moreover, given the extensive set of Cas9-based molecular tools to target activities to DNA sequences (Cong et al., 2013; Kiani et al., 2015; Knott and Doudna, 2018; Konermann et al., 2015; Mali et al., 2013; Perez-Pinera et al., 2013; Sternberg and Doudna, 2015), pGuide cascades should provide new capabilities for engineering mammalian cells and tissues by enabling delivery of gene regulatory activities to sites via preprogrammed sequential orders.

## ACKNOWLEDGEMENTS

We thank Brian Shy for discussions, Kevin Kunstman and Stefan Green in the DNA Sequencing Core, and Balaji Ganesh and Suresh Ramasamy in the Flow Cytomtery core within UIC Research Resources Center for assistance with next generation sequencing and flow cytometry. This work and the personnel were supported by grant funding from the National Institutes of Health (R21 OD027080 to BJM, F30 CA225058 to ART, F30 HD090938 to MSM) and the UIC Center for Clinical and Translational Sciences (RC, HP, MSM)

## AUTHOR CONTRIBUTIONS

Conceptualization, methodology, and writing; RC, BJM.

Experimental execution; RC, ART, HP, MSM, MR.

Data analysis/visualization; RC, ART, BJM

Supervision; BJM.

## AUTHOR INFORMATION

The following competing interests are declared for RC, ART, HP, MSM, MR, and BJM as shareholders in Cellgorithmics, Inc., RC, HP, MSM, and BJM as cofounders of Cellgorithmics, Inc., and RC, HP, MSM, and BJM inventors on patent application number PCT/US2018/052211.

## EXPERIMENTAL PROCEDURES

### Plasmid construction and spacer cloning

Construction of pGuides and cloning of all spacers utilized pSpgRNA as the base plasmid (Addgene, #47108). All sgRNAs were cloned following the protocol optimized for pX330 base plasmids https://www.addgene.org/crispr/zhang/. sgRNA oligo sequences listed without BpiI sticky ends used for cloning and all primers associated with pGuide cloning can be found in Supplementary Table 1.

Generation of pGuide base plasmids lacking ribozymes: In order to facilitate facile cloning of a sequence containing a ribozyme flanked by two Cas9 target sites, base plasmids harboring EcoRI and BamHI consensus sequences within the tetraloop or hairpin 1 were generated using pSpgRNA. The restriction endonuclease sites were flanked by 20 bp that are complementary to each other as a transcript. This 20 bp sequence functions as an insulator to withhold the secondary structure of the tetraloop or hairpin after ribozyme excision. The insulator and restriction site DNA sequence also contained homology to the insertion site within the sgRNA scaffold for Gibson Assembly. The insert was synthesized by IDT and amplified via PCR using Phusion high GC buffer (NEB) supplemented with 3% dimethylsulfoxide (DMSO) and standard PCR conditions (98°C for 30s, 30 cycles of 98°C for 5s, 64°C for 10s and 72°C for 5s, and one cycle of 72°C for 5m). PCR products were column purified (Zymo DNA clean & concentrate). Successful Gibson Assembly reactions occurred using a 7:1 molar ratio of insert:backbone.

Cloning of ribozymes into pSpgRNA to generate pGuides: Ribozyme containing sequences were generated through PCR amplification of the hammerhead ribozyme sequence while simultaneously attaching a 5’ EcoRI site upstream of a Cas9 target site and a 3’ BamHI site downstream of the other Cas9 target site. PCR reactions were performed using Phusion high HF buffer (NEB) and standard PCR conditions (98°C for 30s, 30 cycles of 98°C for 5s, 64°C for 10s and 72°C for 5s, and one cycle of 72°C for 5m). PCR products were column purified (Zymo DNA clean & concentrate) and digested with EcoRI and BamHI alongside digestion of the base plasmid. All digested products were column purified and mixed at a 6:1 molar ratio in the presence of T4 DNA ligase (NEB).

Cloning of pGuides into piggybac transposon vectors: A piggybac transposon vector harboring a puromycin resistance gene (Addgene, #104537) was digested in reactions containing AgeI, NotI and Calf intestinal phosphatase (NEB). In parallel, pGuides located within pSpgRNA were amplified via PCR following standard conditions mentions above. pGuide PCR products were digested with AgeI and NotI, and subsequently mixed at a 6:1 ratio with the digested backbone in reactions containing T4 DNA ligase (NEB).

Generation of constructs encoding Cas9Gem and Cas9Cdt1 fusion proteins: Cdt1 and Geminin degron sequences (Sakaue-Sawano et al., 2008) were ordered as gBlocks (IDT; Extended Data Table 1) containing gly-gly-ser-ser-gly-gly linkers upstream of the degron and flanked by Gibson homology arms for cloning into the carboxy terminus of Cas9. pX330, CyclinE-WTCas9, and CyclinB-WTCas9 were digested with FseI (NEB), followed by mixing with the gBlock DNAs in Gibson Assembly reactions.

### In vitro transcription

Templates for *in vitro* transcription were generated via PCR mediated fusion of the T7 RNAP promoter to the 5’ end of the sgRNA sequence using the appropriate pSPgRNA as the reaction template DNA. PCR reactions were performed using Phusion high HF buffer (NEB) and standard PCR conditions (98°C for 30s, 30 cycles of 98°C for 5s, 64°C for 10s and 72°C for 15s, and one cycle of 72°C for 5m). PCR products were then column purified (Qiagen) and eluted in TE (10mM Tris-HCl pH 8.0, 1mM EDTA). DNA concentrations were determined using a Nanodrop 2000 (ThermoFisher Scientific), and then were diluted to 200nM when used as templates for *in vitro* transcription reactions. The transcription reactions contained 5.0μg/ml purified recombinant T7 RNAP (a gift from Dr. Miljan Simonovic) and 1x transcription buffer (40mM Tris-HCl pH8.0, 2mM spermidine, 10mM MgCl2, 5mM DTT, 2.5mM rNTPs).

Following incubation at 37°C for 1 hour, reactions were treated with RNase free DNase I (ThermoFisher Scientific) and column purified using the Zymo RNA Clean & Concentrator kit following the manufacturer’s protocol. The purified RNA products were eluted from the column in 15μl of water and loaded onto TBE-urea gels for analysis.

### Cell culture

mESC::Rex1-EGFPd2 (A gift from Dr. Austin Smith) were cultured on gelatin in growth medium consisting of N2B27 supplemented with 1 mM PD0325901 (MEKi, Stemgent), 3 mM CHIR99021 (GSK3i, Sigma), and 1000 units/mL LIF (Millipore) (2iL). Cell cultures were routinely split 1:10 with 0.25% trypsin–EDTA every 2–3 days.

HEK293 cells (A gift from Dr. Sojin Shikano), mouse 4T1::EGFP cells (a gift from Dr. Nissim Hay), and HEK293::TLR (a gift from Dr. William Putzbach) were cultured in high glucose DMEM supplemented with 10% FBS and 1% Pen/Strep and were split every 2-3 days using 0.25% trypsin–EDTA.

### Generation of cell lines

Cell lines were generated through transient transfection of piggbac transposase plasmids (HyperPiggybac) and respective pGuide transposon plasmids harboring puromycin resistance cassettes. Briefly, 150,000 4T1 cells were seeded onto 24 wells plates or 225,000 mES cells were seeded onto gelatin coated 24 well plates, followed by transfection of 100 ng transposase plasmid and 400 ng transponon plasmid prepared via mixing with Lipofectamine 2000. As a negative control for selection, transfection mixes containing transposon plasmids only were applied. Media was changed 16 hours after transfection and cells were recovered in respective medias for 48 hours total. To select for integration of the transposon, 2 μg per ml puromycin (Gibco) was applied to mESC for 10 days and 4 μg per ml puromycin was applied to 4T1 for 7 days. Selection was halted once the negative control conditions contained no viable cells.

### TLR and ouabain experiment: transfections, culturing, imaging, and quantification

Transfection: 90,000 HEK293::TLR cells were seeded into each well of a 48 well plate that was pre-coated with poly-l-lysine (Sigma). Immediately after seeding, cells were transfected with guide RNA and Cas9 expression plasmids using Lipofectamine 2000. Briefly, 31.25 ng of each guide RNA expression plasmid was mixed with 125 ng of WTCas9 or Cas9Cdt1 expression plasmids. Each condition contained a total of 250 ng of plasmid DNA. In conditions where the sum of guide RNA and Cas9 plasmid masses did not equal 250 ng, filler plasmid DNA was incorporated to bring the final mass to 250 ng. Plasmid mixes were prepared in Optimem (Gibco) and mixed with 0.75 μl of LF2000 and added dropwise to cells, which incubated with the liposomes for 16 hours before media was changed.

Ouabain selection and culturing: 48 hours and 96 hours after transfection cells were split and the concentration of cells from one representative control well was measured followed by transfer of ~30,000 cells from each condition to black-walled, clear bottom 96 well plates (Greiner) coated with poly-l-lysine. Cells were seeded into media containing ouabain (Tocris) at a final concentration of 0.5 μM. Media containing 0.5 μM ouabain was changed every 2 days for a total of 6 days without splitting before assessing viability.

Imaging-based measurement of red fluorescence and viability: Once the cells settled to the bottom of 96-well black walled plates (~45 minutes after splitting into ouabain), initial cell number and red fluorescence were measured using a Celigo (Nexcelom). Briefly, both red fluorescence and bright field images were acquired. To quantify red fluorescence, “Target 1 + Mask” analysis settings were used which first identifies cells via the bright field image and then quantifies the mean red fluorescence within each cell. This analysis also provides the starting number of cells within each well. After 6 days of culturing in ouabain, viability was assessed via staining for all cells using Hoeschst33342 (ThermoFisher) and live cells using Calcein AM (Life Technologies). Briefly, media containing 10 μM Hoeschst and 1 μM Calcien AM was added to every condition and incubated for 1 hour at 37°C in the dark. Subsequently, every condition was imaged using the Celigo, and relative viability was calculated by dividing the number of Hoechst positive/Calcein AM positive cells by the number of cells initially seeded in each well, as measured by the Celigo 6 days prior during seeding.

### Western blot analysis

~2 million ES cells were collected from 6 well plates after washing twice with PBS, then scraping into 600ul of PBS followed by pelleting at 5,000 rpm for 5 minutes. The cell pellet was then resuspended in 100ul of pre-heated (98°C) 2x Laemmli lysis buffer (4% SDS, 20% glycerol, 120mM Tris-Cl, 0.02% w/v bromophenol blue) and heated at 98°C for 10m. While still hot, each sample was resuspended with a 25-gauge needle to shear genomic DNA. 25μl of each sample was loaded onto a 12% SDS PAGE gel and then transferred to 0.45 mm nitrocellulose membranes (Bio-rad). Protein loading was assessed through ponceau staining of nitrocellulose membranes. Membranes were blocked for 1 hour at room temperature with 5% milk – TBST (Tris buffered saline, 0.1% tween) before probing for anti-FLAG (Sigma). The primary antibody was diluted 1:2000 in 5% milk - TBST and was incubated at room temperature for 1 hour, followed by 3 washes with TBST. An anti-mouse secondary antibody conjugated to horse radish peroxidase (HRP) (Jackson Labs) was diluted 1:5000 in 5% milk – TBST, applied, then incubated for 1 hour at room temperature.

### Flow cytometry

Single-cell suspensions were prepared by trypsinization and re-suspension in 2% FBS/PBS/2mM EDTA, followed by filtering through a 35 μm strainer cap. All data was collected on a CytoFLEX S flow cytometer (Beckman Coulter), and data analysis was performed using CytExpert software (Beckman Coulter). To isolate live and single cells, gating by forward scatter and side scatter area was applied followed by side scatter area and side scatter width. At least 5 × 10^5^ singlet, live cells were counted for each sample. For transient transfections measuring EGFP disruption, mCherry positive live singlet cells were gated and EGFP positive and negative populations were measured.

### Next generation DNA sequencing

Genomic DNA was harvested via trypsinization and pelleting of cells, followed by resuspension in Quick Extract (Lucigen). Samples were vortexed at top speed for 15 seconds, incubated at 65°C for 6 minutes, vortexed for 15 seconds, incubated at 98°C for 2 minutes, then vortexed for 15 seconds. Approximately 100ng of DNA was used in PCR to amplify respective target sites while attaching adapter sequences (Fluidigm) for subsequent barcoding steps (Extended Data Table 1 for NGS primers). PCR products were analyzed via agarose gel and then distinct amplicons were pooled for each replicate respectively in equal amounts based on ImageJ quantification. Pooled PCR products were purified with AMPure beads (Agilent), and 5ng of the purified pools was barcoded with Fluidigm Access Array barcodes using AccuPrime II (ThermoFisher Scientific) PCR mix (95°C for 5m, 8 cycles of 95°C for 30s, 60°C for 30s and 72°C for 30s, and one cycle of 72°C for 7m). Barcoded PCR products were analyzed on a 2200 TapeStation (Agilent) before and after 2 rounds of 0.6x SPRI bead purification to exclude primer dimers. A final pool of amplicons was created and loaded onto an Illumina MiniSeq generating 150bp paired-end reads.

### Next generation DNA sequencing analysis

Determination of indel frequencies made use of CRISPResso version 2 command line tools that demultiplexed by amplicon (Clement et al., 2018), where appropriate, and then determined indel frequency by alignment to reference amplicons. Outputs were assembled and analyzed using custom command-line, python, and R scripts which are available upon request.

### Nucleic acid sequences

Tetraloop base sequence with EcoRI and BamHI sites:

GTTTTAGAGCTAACGTGCGATCGGTCTGACGTgaattcTATAGggatccACGTCAGACCGATCGCACGTTAGCAAGTT AAAATAAGGCTAGTCCGTTATCAACTTAAGTGGCACCGAGTCGGTGCTTTT

Hairpin 1 base sequence with EcoRI and BamHI sites:

GTTTTAGAGCTAGAAATAGCAAGTTAAAATAAGGCTAGTCCGTTATCAACTTACGGGCGGTCGGCCTGGCGTgaatt cTATAGggatccACGCCAGGCCGACCGCCCGTAAGTGGCACCGAGTCGGTGCTTT

Twister ribozyme: GGTGCCTAACACTGCCAATGCCGGTCCCAAGCCCGGATAAAAGTGGAGGGGGCA

Hepatitis delta ribozyme: GGGGTGCTTCGGATGCTGATGAGTCCGTGAGGACGAAACAGGGCAACCTGTCCATCCGGTATCCC

Hammerhead ribozyme: CCTGTCACCGGATGTGCTTTCCGGTCTGATGAGTCCGTGAGGACGAAACAGG

Stem 1 pGuide:

GTTTTAGAGCTAACGTGCGATCGGTCTGACGTgaattcTAGCGCTCCTACGGAAGTCGGGAAGGtcgCCTGTCACCG GATGTGCTTTCCGGTCTGATGAGTCCGTGAGGACGAAACAGGcgattttttGCGCTCCTACGGAAGTCGGGAAGGTA GggatccACGTCAGACCGATCGCACGTTAGCAAGTTAAAATAAGGCTAGTCCGTTATCAACTTAAGTGGCACCGAGT CGGTGCTTTT

Stem 2 pGuide:

GTTTTAGAGCTAACGTGCGATCGGTCTGACGTgaattcTACCTTCCCGACTTCCGTAGGAGCGCtcgCCTGTCACCGG ATGTGCTTTCCGGTCTGATGAGTCCGTGAGGACGAAACAGGcgattttttCCTTCCCGACTTCCGTAGGAGCGCTAGgg atccACGTCAGACCGATCGCACGTTAGCAAGTTAAAATAAGGCTAGTCCGTTATCAACTTAAGTGGCACCGAGTCG GTGCTTTT

Stem 3 pGuide:

GTTTTAGAGCTAACGTGCGATCGGTCTGACGTgaattcTAGTCAAGTAGACGGTACAACGACGGtcgCCTGTCACCG GATGTGCTTTCCGGTCTGATGAGTCCGTGAGGACGAAACAGGcgattttttGTCAAGTAGACGGTACAACGACGGTA GggatccACGTCAGACCGATCGCACGTTAGCAAGTTAAAATAAGGCTAGTCCGTTATCAACTTAAGTGGCACCGAGT CGGTGCTTTT

Stem 4 pGuide:

GTTTTAGAGCTAACGTGCGATCGGTCTGACGTgaattcTACCGTCGTTGTACCGTCTACTTGACtcgCCTGTCACCGGA TGTGCTTTCCGGTCTGATGAGTCCGTGAGGACGAAACAGGcgattttttCCGTCGTTGTACCGTCTACTTGACTAGggat ccACGTCAGACCGATCGCACGTTAGCAAGTTAAAATAAGGCTAGTCCGTTATCAACTTAAGTGGCACCGAGTCGGT GCTTTT

Stem 5 pGuide:

GTTTTAGAGCTAACGTGCGATCGGTCTGACGTgaattcTAGAGGACCCGATCTTTACCTTCAGGtcgCCTGTCACCGG ATGTGCTTTCCGGTCTGATGAGTCCGTGAGGACGAAACAGGcgattttttGAGGACCCGATCTTTACCTTCAGGTAGgg atccACGTCAGACCGATCGCACGTTAGCAAGTTAAAATAAGGCTAGTCCGTTATCAACTTAAGTGGCACCGAGTCG GTGCTTTT

Stem 6 pGuide:

GTTTTAGAGCTAACGTGCGATCGGTCTGACGTgaattcTACCTGAAGGTAAAGATCGGGTCCTCtcgCCTGTCACCGG ATGTGCTTTCCGGTCTGATGAGTCCGTGAGGACGAAACAGGcgattttttCCTGAAGGTAAAGATCGGGTCCTCTAGg gatccACGTCAGACCGATCGCACGTTAGCAAGTTAAAATAAGGCTAGTCCGTTATCAACTTAAGTGGCACCGAGTCG GTGCTTTT

Cdt1 degron:

CCTTCCCCTGCTAGGCCTGCTCTGCGCGCTCCAGCTTCTGCCACATCTGGGAGTCGAAAACGCGCTCGACCTCCCGC CGCTCCGGGACGCGACCAGGCCAGGCCTCCCGCCAGACGGCGGCTGCGACTGTCCGTGGACGAGGTGAGCTCAC CCTCCACACCTGAAGCCCCTGATATCCCCGCTTGTCCATCTCCTGGGCAGAAAATCAAAAAGAGCACCCCGGCTGC AGGGCAGCCTCCACACCTTACTTCTGCCCAAGATCAGGATACCATCTAA

Gem degron:

ATGAATCCCAGTATGAAGCAGAAACAAGAAGAAATCAAAGAGAATATAAAGAATAGTTCTGTCCCAAGAAGAACT CTGAAGATGATTCAGCCTTCTGCATCTGGATCTCTTGTTGGAAGAGAAAATGAGCTGTCCGCAGGCTTGTCCAAAA GGAAACATCGGAATGACCACTTAACATCTACAACTTCCAGCCCTGGGGTTATTGTCCCAGAATCTAGTGAAAATAA AAATCTTGGAGGAGTCACCCAGGAGTCATTTGATCTTATGATTAAAGAAAATCCATCCTCTCAGTATTGGAAGGAA GTGGCAGAAAAACGGAGAAAGGCGCTGTAA

GGSSGG linker:

GGGAGTGGCGGTTCTGGA

---

**Supplementary Figure S1.**
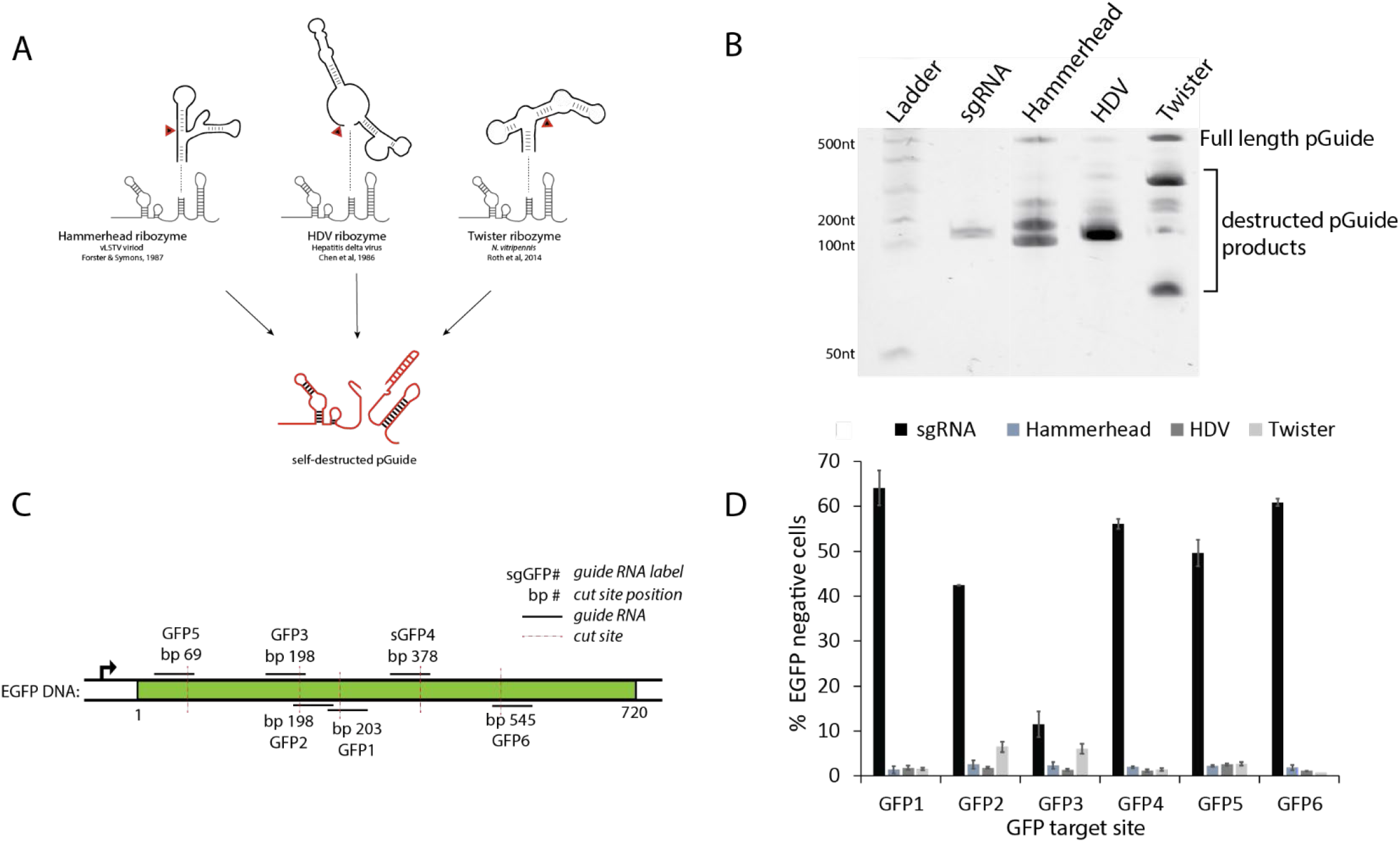
Related to Figure 1 and 2: pGuide inactivity is RNA-based on *cis*-cleaving ribozymes. A. Several modalities for inactivating sgRNA were tested via insertion of elements into the sgRNA-encoding gene. Initial testing of ribozymes used insertions of a new sequence at the end of hairpin 1. Shown here are the structures of three ribozymes that were inserted and tested *in vitro* and in cells. The red arrow denotes the cis-cleavage site for each ribozyme. B. pGuides self-destruct during *in vitro* transcription. Each pGuide variant containing a unique ribozyme at hairpin 1 was cloned into plasmid in which a T7 promoter directed *in vitro* transcription of the pGuide. Transcription reaction products were analyzed via EtBr staining on a TBE-urea denaturing gel. The sgRNA gene (no ribozyme insertion) generated a single 100nt band, corresponding to intact sgRNA. All three pGuide gene variants result in high levels of cleavage of the transcribed RNA. Low levels of full-length, uncleaved pGuide were at detected at varying amounts depending on the ribozyme. The HDV ribozyme appeared to be the most efficient *in vitro*. Uncleaved ribozymes may result from an insufficient levels of Mg^2+^ in the *in vitro* reaction; Mg^2+^ is used both by T7 RNA polymerase for transcription and by ribozymes as a cofactor required for catalysis. C. Schematic showing locations of target sites utilized for assessing genome editing by pGuides. The DNA sequences for sites GFP1-6 are listed in Table S1. Base pair numbering begins with +1 at the first coding nucleotide for EGFP. D. Mouse 4T1 cells harboring a single genomic insertion of a heterologous EGFP gene were used to assess spCas9 activity. In separate transient transfections, sgRNA targeting six different sites (sgGFP1-6) throughout the EGFP coding sequence were tested. For each target site, the normal sgRNA without a ribozyme insertion and pGuides with either a Hammerhead, HDV, or Twister ribozyme were generated and tested. Mutagenesis of the EGFP gene is inferred from loss of EGFP fluorescence measured by flow cytometry performed five days after transfections. Data represent mean +/−range of n = 2 biological replicates.

**Supplementary Figure S2.**
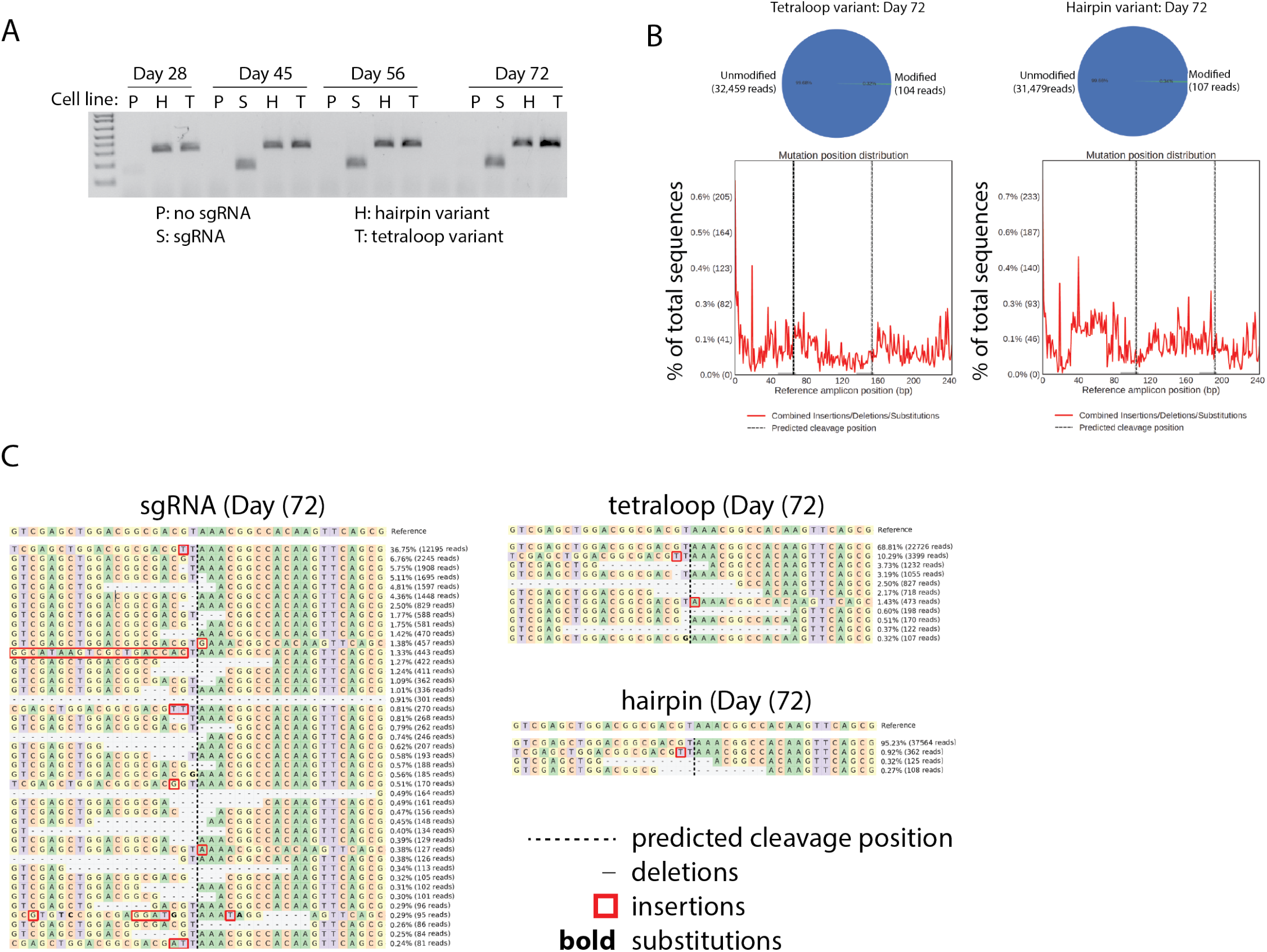
Related to Figure 2 and 3: pGuide inactivity persists over long timespans while maintaining competence for conversion to functional mGuides. A,B. Stability of pGuide genes integrated into 4T1 cells was assessed by PCR amplification of the pGuide sequence or sgRNA control sequences. 4T1 cell lines expressing spCas9 from a lentiviral insertion and either one of the two pGuide variants, an sgRNA, or no sgRNA were generated using piggyBac transposase. Cells were continuously expanded in culture (Fig. 2D) and genomic DNA was isolated from cells at the indicated days. PCR products were examined by agarose gel electrophoreses (A) and targeted deep sequencing (B). In (B), pie charts (above) show the number of unmodified sequencing reads versus modified sequencing reads for the tetraloop and hairpin variants. Analysis of modified reads for each sample (below) shows a sequencing distribution pattern that resembles sequencing noise at background levels. Location of the ribozyme is demarcated by the two dashed lines. Data is representative, from a polyclonal population of ~1,000,000 cells. C. Representative allele maps depicting the abundance of EGFP mutant alleles generated by the sgRNA, tetraloop pGuide or hairpin pGuide after 72 days of continuous culturing. Allele maps were obtained using CRISPResso2-based analysis of sequencing data associated with Fig. 2D. Alleles are ordered from high abundance (top) to low abundance (bottom). The alleles presented in each map do not represent all mutant alleles associated with each NGS sample.

**Supplementary Figure S3.**
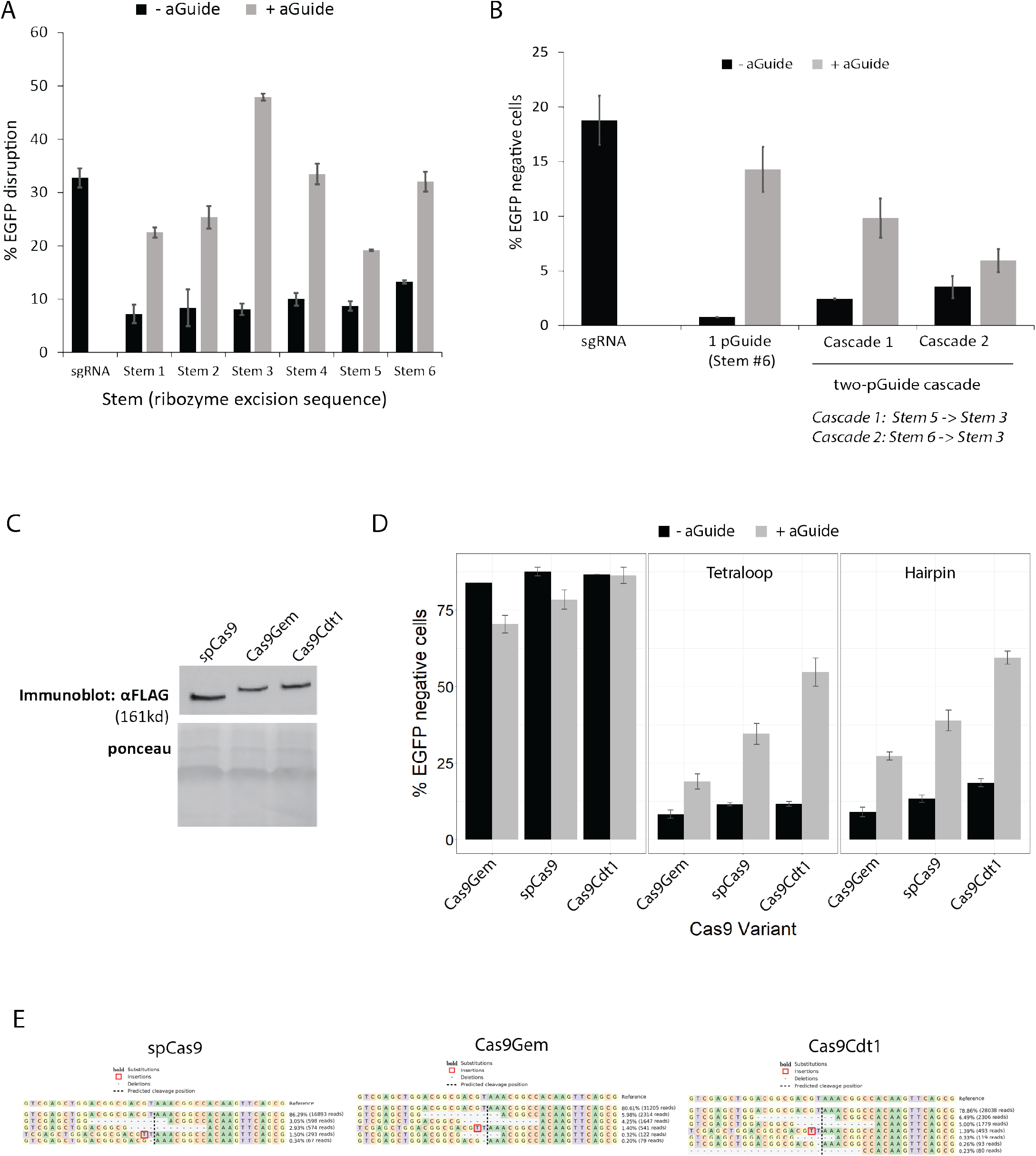
Related to Figures 4-6: Cell cycle regulation of spCas9 modulates mGuide mutagenesis kinetics. A. Data show raw values prior to normalization corresponding to Fig. 4D. The effect of the stem sequence (Cas9 target sites flanking the ribozyme) on pGuide to mGuide conversion was examined through generating 6 different tetraloop pGuide variants harboring different stem sequences, followed by an EGFP disruption assay to measure Cas9 activity. mESC::Rex1 cells were transfected with guide RNA and Cas9 expression plasmids, followed by flow cytometry analysis of EGFP levels after 48 hours. Data are displayed as mean +/−range of n = 2 biological replicates. Stem nucleotide sequences are located in Table S1. B. Data represents raw values prior to normalization corresponding to Fig. 5B. A linear cascade of two pGuides executing three sequential genome edits was assessed. Guide RNA plasmids were transfected into cells and EGFP levels were measured 48 hours after via flow cytometry. Data are displayed as standard deviation of n = 3 biological replicates. C. Expression of Cas9-degron fusions. Protein levels of the spCas9 and variants were measured by western blot 24 hours after transiently transfecting Rex1::GFPd2 cells with each plasmid. Cas9Gem and Cas9Cdt1 exhibit ~50% less protein levels than spCas9 and migrate at a higher molecular weight due to fusion of the degron to Cas9. D. Activation of episomal pGuides with Cas9-degron fusions reveals different kinetics associated with cell cycle regulation of Cas9. Data are displayed as mean +/−standard deviation of n = 3 biological replicates. E. Next generation sequencing confirms that modulation of pGuide activation via Cas9Cdt1 and Cas9Gem is due to increases or decreases in mutation frequencies, not changes in mutational outcomes. We examined the possibility that EGFP disruption frequencies were different as a result from cell cycle timed DSBs altering the frequency of in frame mutations relative to frameshift indels. Allele frequencies were determined by deep sequencing of DNA harvested from Rex1::GFPd2 cells 48 hours after transfection with tetraloop pGuides and the spCas9 or variant plasmids from experiment shown in (D). Analysis reveals three informative results: 1) the mutational profiles generated by spCas9 and the two variants are highly concordant, 2) mutation frequencies are increased or decreased for Cas9Cdt1 and Cas9Gem, respectively, and 3) the top two mutant alleles are in frame deletions (12 bp), potentially leading to an observed EGFP loss values that are below actual mutation frequencies. Allele maps are representative of one of two replicates.

**Supplementary Figure S4.**
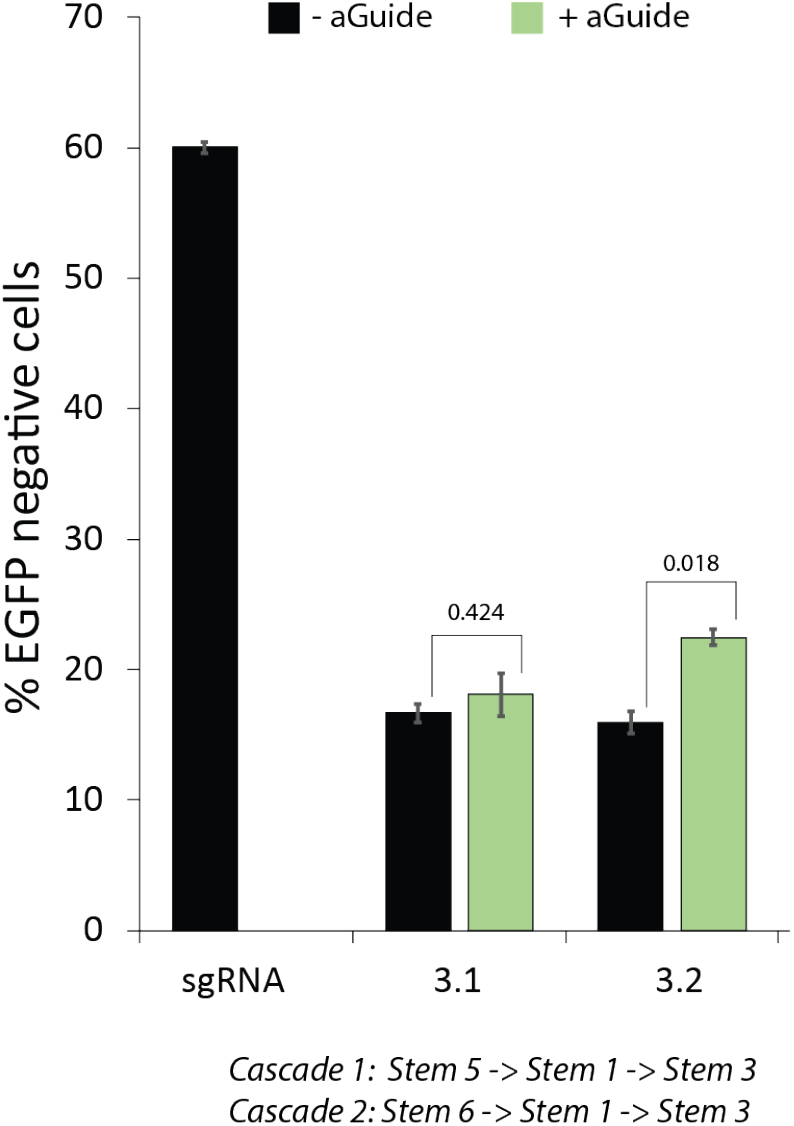
Related to Figure 5: Four event linear cascade of pGuides in cells. Transient transfection of guide RNA and spCas9 expression plasmids into Rex1::GFPd2 cells followed by analysis of EGFP levels was performed to assess the efficiency of cascade completion. Experiment and analyses are identical to those for Fig 5F, with the exception that spCas9 was used instead of Cas9Cdt1. Error bars represent +/−s.d. for n = 3 biological replicates. p-values were calculated using Welch’s two sided t-test.

**Supplementary Figure S5.**
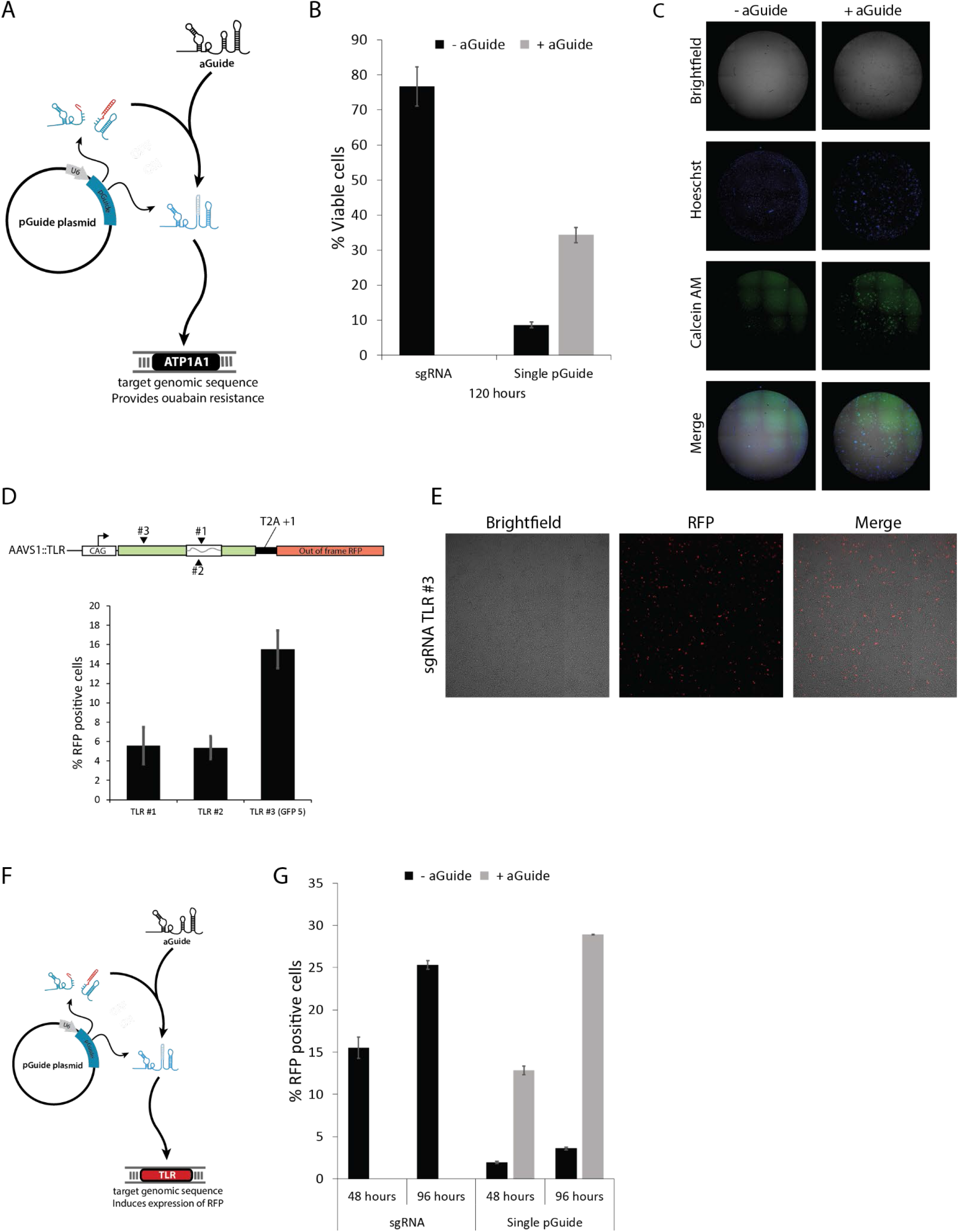
Related to Figure 6: Functionality of individual arms of the ramified cascade of pGuides. A. Schema depicting single pGuide activation that results in mutagenesis of APT1A1 and resistance to ouabain. B. A previously reported sgRNA sequence (REF)(Agudelo et al, *Nat. Meth.* 2017) was used to generate ouabain-resistance mutations in Cas9-expressing cells. The same target sequence was used for a pGuide, which was activated in cells with co-expression of an aGuide. The ability of sgRNA or pGuide to mediate ouabain resistance was compared by transfecting HEK293 cells with guide RNA and Cas9 expression plasmids. Cells were by treated with ouabain five days after transfection and allowed to grow for six days before measuring viability. Viability was determined by staining cells with Calcein AM and dividing the number of Calcein AM positive cells by the total number of starting cells for each condition. Data represent mean +/−standard deviation of n = 3 biological replicates. C. Representative images from the Celigo detecting all cells: brightfield and Hoeschst, and live cells: Calcein AM in conditions containing pGuides only (-aGuide) or mGuides (+ aGuide). D. For the traffic light reporter gene to be converted to encode for a functional red fluorescent protein (RFP), a +2bp insertion or a - 1bp deletion is required to shift the frame and enable the RFP to be translated in frame. (Top). Three sgRNA (TLR#1-#3) were assessed for the frequency by which they generated RFP-positive cells after transfection into AAVS::TLR cells. Note that TLR#3 is identical to GFP5 (Fig. S1C). (bottom) RFP positive cells were detected using Celigo-based imaging. Brightfield was used as the measure for total cell number and the subsequent denominator to determine percent RFP positive cells. Data are displayed as the mean +/−standard deviation of n = 3 biological replicates E. Representative images from the Celigo instrument used for counting total cells and RFP-positive cells and used for comparison of sgRNA in panel A. F. Schematic depicting activation of a single pGuide to target the TLR and induce expression of RFP. G. Induction of RFP resulting from sgRNA or a tetraloop pGuide was measured at 48 and 96 hours post transfection of sgRNA and Cas9 expression plasmids. Data represent mean +/−standard deviation of 3 biological replicates.

**Supplementary Figure S6.**
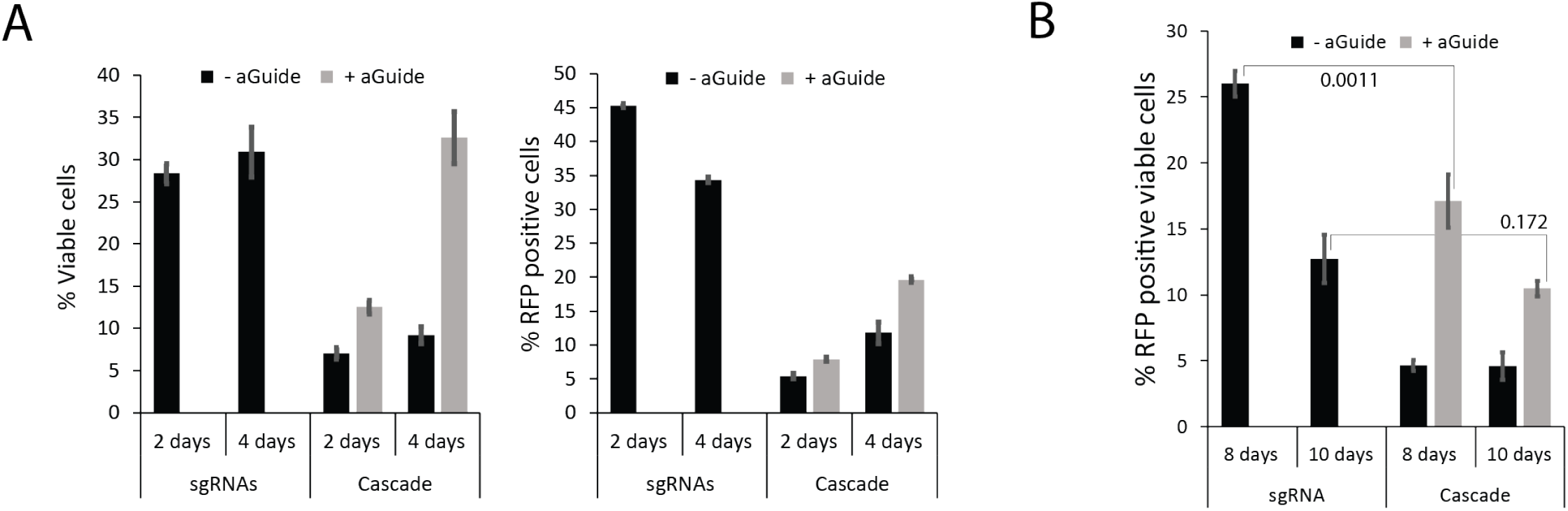
Related to Figure 6: A. Data represents raw values prior to normalization corresponding to Fig. 6B. Plasmid DNAs encoding Cas9Cdt1 and pGuides targeted to ATP1A1 and the TLR were transfected into HEK293 harboring a single genomic TLR cassette (see Fig. 6A for sequential targeting strategy). Two and four days after transfections, cells were plated into ouabain followed by immediate measurement of RFP positive cells using a Celigo imager. After 6 days of continuous culturing in ouabain, viable cells were measured via Calcein AM staining and divided by the initial cell number measured upon plating into ouabain. Data represent mean +/−standard deviation of 3 biological replicates. B. Data represents raw values prior to normalization corresponding to Fig. 6D. To compare the efficiency and kinetics of the pGuide cascade to sgRNA in generating double positive ouabain resistant, RFP expressing cells, viable cells that were all Calcein AM positive cells were counted followed by assessment of RFP positivity within the viable subpopulation. p-values are presented to compare the sgRNA condition to the pGuide condition to address differences in kinetics, revealing that the pGuide cascade exhibits delayed kinetics but similar efficiencies as sgRNAs. p-values are located above or below the lines drawn between two conditions and were calculated using Welch’s two sided t-test. Data represent mean +/−standard deviation of n=3 biological replicates.

